# Improving the prediction of protein stability changes upon mutations by geometric learning and a pre-training strategy

**DOI:** 10.1101/2023.05.28.542668

**Authors:** Yunxin Xu, Di Liu, Haipeng Gong

## Abstract

Accurate prediction of the fitness and stability of a protein upon mutations is of high importance in protein engineering and design. Despite the rapid development of deep learning techniques and accumulation of experimental data, the multi-labeled nature of fitness data hinders the training of robust deep-learning-based models for the fitness and stability prediction tasks. Here, we propose three geometric-learning-based models, GeoFitness, GeoDDG and GeoDTm, for the prediction of the fitness score, ΔΔ*G* and Δ*T*_*m*_ of a protein upon mutations, respectively. In the optimization of GeoFitness, we designed a novel loss function to allow supervised training of a unified model using the large amount of multi-labeled fitness data in the deep mutational scanning (DMS) database. By this means, GeoFitness efficiently learns the general functional effects of protein mutations and achieves better performance over the other state-of-the-art methods. To further improve the downstream tasks of ΔΔ*G*/Δ*T*_*m*_ prediction, we re-utilized the encoder of GeoFitness as a pre-trained module in GeoDDG and GeoDTm to overcome the challenge of lack of sufficient amount of specifically labeled data. This pre-training strategy in combination with data expansion remarkably improves model performance and generalizability. When evaluated on the benchmark test sets (S669 for ΔΔ*G* prediction and a newly collected set S571 for Δ*T*_*m*_ prediction), GeoDDG and GeoDTm outperform the other state-of-the-art methods by at least 30% and 70%, respectively, in terms of the Spearman correlation coefficient between predicted and experimental values. An online server for the suite of these three predictors, GeoStab-suite, is available at http://structpred.life.tsinghua.edu.cn/server_geostab.html.

## Introduction

Protein fitness is defined as the ability of a protein to perform a specific function, but is frequently quantified by different indicators (*e.g*., enzyme activity, peptide binding affinity and protein stability)^1^ in various experimental cases. One of the main goals of protein design and engineering is to improve the protein fitness so as to enhance protein performance in biotechnological and biopharmaceutical processes^2,3^. Among the various indicators of protein fitness, the protein stability^4^ is of high interest, which is commonly evaluated by two metrics, Δ*G* and *T*_*m*_. Δ*G* denotes the unfolding free energy change at room temperature and describes the thermodynamic stability of a protein, while *T*_*m*_ stands for the protein melting temperature and reflects the ability of a protein to maintain its folded state against temperature fluctuations^5,6^. Understanding protein stability not only facilitates the acquisition of engineered proteins with sustained activities in harsh bioprocesses and/or under manufacturing conditions^7^, but also aids in the elucidation of the role of human genetic variants in the etiology of diverse diseases in the field of biomedicine^8^. Indeed, much attention has been paid to learn the changes of fitness (and particularly the stability) of proteins upon mutations. For instance, the deep mutational scanning (DMS) database^9^ contains over 3,000,000 pieces of protein fitness data, which were measured using various experimental indicators upon mutations. The experimental strategy is, however, highly costly and labor-intensive, considering the excessively large number of possible protein sequences even for single-point mutations^10–12^. Therefore, developing computational tools for fast and accurate prediction of the mutation effects on protein fitness is in urgent demand.

The protein fitness prediction methods can be developed and optimized based on the DMS database. However, the multi-labeled nature of DMS data hinders the training of a unified prediction model (Figure 1). Existing models^1,3,4,12–30^ mainly take two strategies to circumvent this obstacle. One strategy taken by methods like ECNet^21^ and SESNet^20^ is to train a specific model based on each definition of fitness in the concerned issue. Therefore, additional fitness data with the same corresponding meaning to the interested protein property are required to retrain these models before the practical prediction. This strategy leads to satisfying performance but sacrifices the model usability. Another strategy is to train the model with the masked language modeling objectives, like in RF_joint_^27,28^, MSA transformer^23^, ESM-1b^29^, ESM-1v^22^ and ESM-2^30^, which completely bypasses the need of any fitness data. This strategy can produce unified models for fast prediction but at the cost of model performance. Presently, there still lacks a robust method to generate a unified fitness prediction model with satisfying performance.

**Figure 1.**
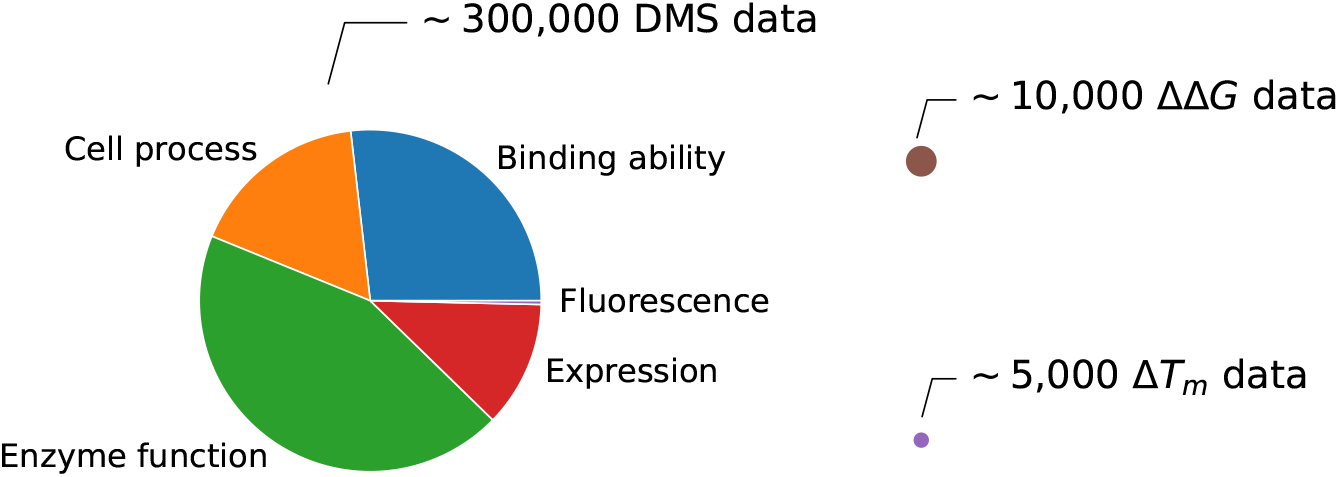
Summary on the DMS, ΔΔ*G*, and Δ*T*_*m*_ data. DMS data are multi-labeled and could be roughly divided into 5 categories. More detailed classification of DMS data utilized in the model training of this work could be found in Table S1. Unlike the DMS data, ΔΔ*G*, and Δ*T*_*m*_ data are single-labeled but have significantly reduced scales, with their data sizes approximately indicated by the radii of circles. In general, although the DMS data adopted in this work have no overlap with the ΔΔ*G*/Δ*T*_*m*_ data, the size of the former is about 20 times of the size of latter ones in combination.

Different from the multi-labeled fitness data, the protein stability changes upon mutations are clearly defined by two metrics, ΔΔ*G* and Δ*T*_*m*_, and the accumulation of experimental data allows the development of corresponding prediction algorithms. Recently, great efforts have been exerted in ΔΔ*G* prediction^31–50^. Current methods can be mainly classified as mechanistic predictors, machine learning predictors, and deep learning predictors^51^. Mechanistic predictors are based on protein energy or statistical potentials, like DDGun and DDGun3D^36^, while machine learning predictors, including INPS-Seq^47^, I-Mutant3.0-Seq^48^, PremPS^49^, *etc*., feed features extracted from protein sequences or structures into various machine learning models for prediction. With the rapid development of deep learning techniques, deep learning models have shown great potential in the accurate prediction of protein stability, exemplified by ACDC-NN-Seq^46^, ACDC-NN^43^ and ThermoNet^45^. Unfortunately, these methods have not demonstrated absolute advantages over traditional methods when evaluated on a newly released benchmark dataset, S669^52^. The critical issue lies in the limited amount of training data (∼ 10, 000), which easily leads to overfitting for deep learning models with millions of parameters. Moreover, available experimental data is biased toward several hotspot protein families, specific target amino acid mutations and destabilizing mutations^6,53^. One manifestation of such biases is that most current predictors are unable to reproduce the anti-symmetry property of ΔΔ*G, i.e*. inversion of the value upon the swapping of the statuses of mutant and wild-type^53^.

Δ*T*_*m*_ prediction is relatively less explored when compared with ΔΔ*G* prediction. Existing models^10,54–59^, like AUTO-MUTE^55,59^ and HoTMuSiC^56^, utilize physicochemical properties or statistical potentials as features and integrate with machine-learning-based methods for prediction. To our knowledge, deep-learning-based methods are absent in this field, possibly due to the significantly reduced amount of available Δ*T*_*m*_ data in comparison with ΔΔ*G*, despite the higher potential impact in practice for the prediction of Δ*T*_*m*_^6^. Besides the same challenges confronted in the ΔΔ*G* prediction, another dilemma in the Δ*T*_*m*_ prediction arises from the lack of established open-source benchmark datasets, which hinders the fair comparison of model performance.

In this work, we first developed a geometric-learning-based model, GeoFitness, for the prediction of protein fitness. Specifically, we designed a novel loss function to allow the training of a unified model with the multi-labeled fitness data in the DMS database. The model derived by this means avoids the prior limitation of model retraining before practical usage and simultaneously achieves improved performance over the other state-of-the-art methods like ECNet. Furthermore, by reutilizing the geometric encoder of GeoFitness, we developed two additional downstream models, GeoDDG and GeoDTm, to predict ΔΔ*G* and Δ*T*_*m*_ of a protein upon mutations, respectively, with the model architecture specifically designed to ensure the anti-symmetry of prediction results. Notably, we addressed the challenge of limited data in ΔΔ*G* and Δ*T*_*m*_ predictions by two strategies: expanding the training data by data collection as well as inheriting the geometric encoder of the GeoFitness model pre-trained on the DMS database. Particularly, the latter strategy remarkably improved the model performance and generalizability, considering the fact that fitness data of protein variants outnumber those of ΔΔ*G* and Δ*T*_*m*_ by at least one order of magnitude as well as the relevance between protein fitness and stability in biology (Figure 1). When evaluated on the benchmark test sets, S669^52^ for ΔΔ*G* and S571 (a newly collected set in this work) for Δ*T*_*m*_ predictions, GeoDDG and GeoDTm outperform the other state-of-the-art methods by at least 30% and 70%, respectively, in terms of the Spearman correlation coefficient between predicted and experimental values.

## Methods

### Overview of the prediction models

The GeoStab-suite developed in this work comprises three distinct software programs, namely GeoFitness, GeoDDG, and GeoDTm, all of which aggregate the information from both protein sequence and structure into a geometric-learning-based encoder for prediction. The Geometric Encoder adopts a graph attention (GAT) neural network architecture, where the nodes (1D) represent amino acid residues and edges (2D) reflect inter-residue interactions (Figure 2a). GeoFitness is a unified model capable of predicting fitness landscape of protein variants of single mutations (Figure 2b). GeoDDG and GeoDTm reutilize the pre-trained information extractor of GeoFitness to predict the ΔΔ*G* and Δ*T*_*m*_ values of proteins upon arbitrary mutations, respectively (Figure 2c). The protein structure information could be either derived from the Protein Data Bank (PDB) or predicted purely based on sequence using AlphaFold2^60^. Consequently, we have trained two versions of GeoDDG and GeoDTm, with the suffixes of “-3D” and “-Seq” to annotate the version that relies on experimental structures and that only needs sequence information in practical usage, respectively.

**Figure 2.**
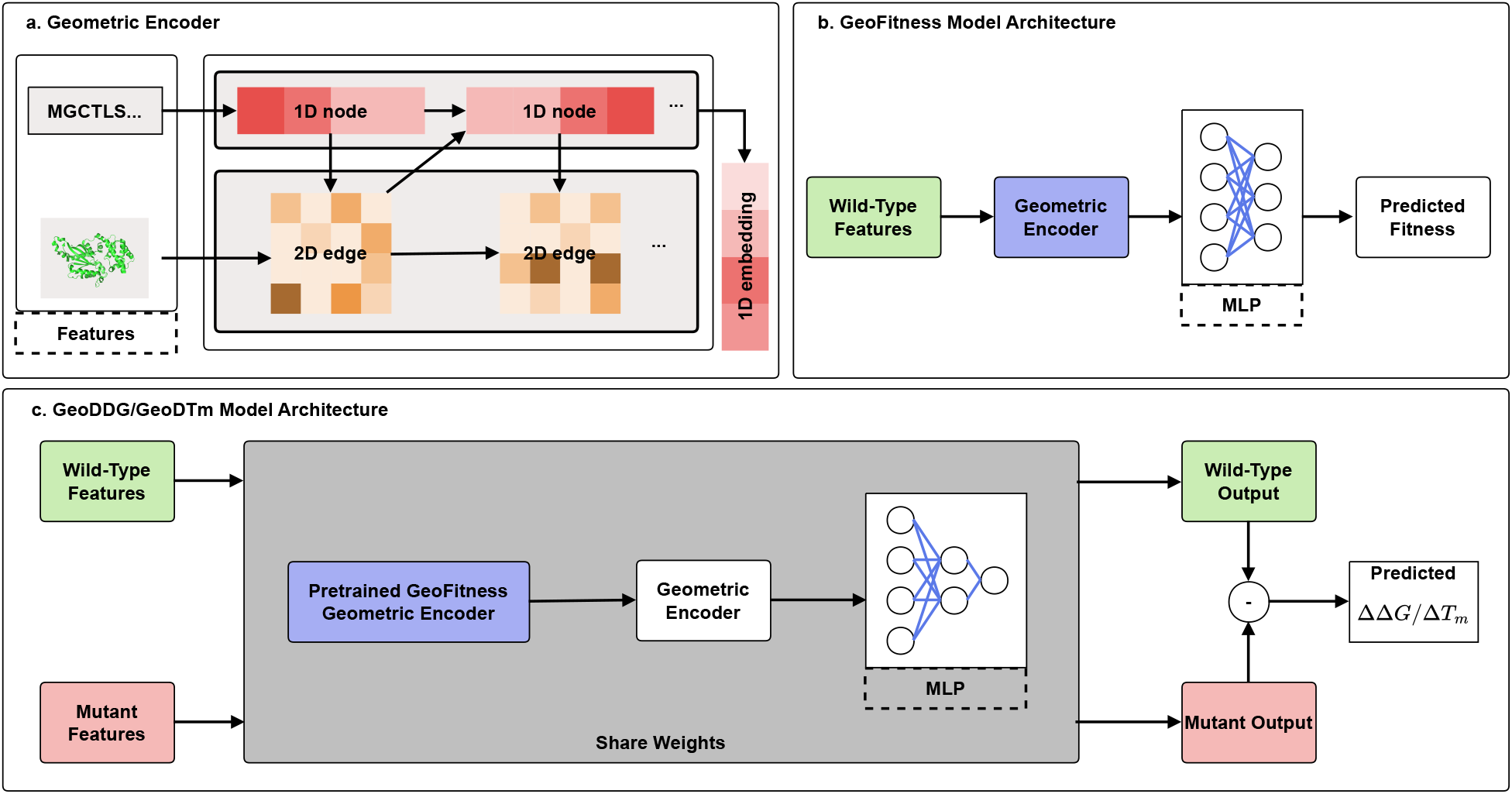
Schematic overview of the model architecture. **a)** The input of Geometric Encoder consists of both sequence and structural information, which initialize the node and edge embeddings, respectively. After alternate updating of nodes and edges in an *N* -layer GAT network, the final embedding is generated. **b)** The GeoFitness model takes the features of wild-type protein sequence and structure as input, and outputs the predicted fitness value through the Geometric Encoder and multi-layer-perceptron (MLP) layers. **c)** The model architectures of GeoDDG and GeoDTm are similiar. Features of wild-type and mutant are processed using the same encoder (with shared weights) and their differences are taken as the prediction results for ΔΔ*G*/Δ*T*_*m*_. Notably, the encoder reutilizes the pre-trained network module of GeoFitness to facilitate information processing, by inheriting the network architecture and parameter values: the modules colored violet in (**b**) and (**c**) are identical.

### Protein fitness database and dataset

#### MaveDB

MaveDB is an open-source database designed to store data of large-scale mutation effects, which was first published in 2019 and had been updated to version 2 in 2021^61,62^. MaveDB includes over 300 datasets related to protein fitness, from which we chose 52 DMS studies on single-point mutation effects. The details of the datasets are summarized in Table S1.

### DeepSequence Dataset

We also filtered DMS data from the DeepSequence dataset^1^ by collecting single-point mutation data but excluding the portions redundant with MaveDB. In this way, we obtained 22 DMS datasets. The details of the datasets are summarized in Table S1.

### Division of training, validation and test sets

The DMS data used in this work contain approximately 300,000 entries from 74 proteins in total (Table S1). For each protein, all available DMS data were randomly assigned as training, validation and testing with a ratio of 7:1:2. In order to reproduce the results of ECNet and SESNet, a specific model was optimized using the corresponding training data for each individual protein. The training data of all proteins were further combined as a comprehensive training set for GeoFitness. Notably, the DMS datasets used here have no overlap in labels with the datasets of protein stability, particularly with the benchmark test sets of ΔΔ*G*/Δ*T*_*m*_ (see Figure S1). Consequently, model pre-training using the plethora of DMS data brings no information leak to the downstream prediction tasks of ΔΔ*G* and Δ*T*_*m*_ as well as the performance evaluation.

### Protein stability database and dataset

#### ProThermDB

ProThermDB, a thermodynamic database for protein mutants, is an upgraded version of its predecessor ProTherm database that was first released in 1999 and then continuously updated until 2006^63–68^. As a refined version, ProThermDB^69^ contains approximately 31,500 pieces of protein stability data with annotation errors in ProTherm corrected.

#### ThermoMutDB

ThermoMutDB is a manually curated thermodynamic database for protein mutants, consisting of two parts of data. The first part is derived from the 1,902 literatures related to the ProTherm database and all data have been double-checked and validated. The second part contains new thermodynamic mutation data collected from approximately 34,000 literatures. The database contains about 15,000 pieces of protein stability data^70^.

#### S2648

The S2648 dataset^33^ is a ΔΔ*G* dataset consisting of 2,648 single-point mutations of 131 proteins collected from ProTherm database. It has been widely utilized as a training set in prior softwares for ΔΔ*G* prediction and is considered the most extensively employed dataset of its kind.

#### S669

The S669 dataset^52^ is made up with 669 single-point mutations of 94 proteins, which have < 25% sequence similarity with those in S2648 and VariBench^71^. As many softwares for ΔΔ*G* prediction have previously utilized S2648 or VariBench for model training, S669 is generally considered as a fair benchmark test set for the performance evaluation of ΔΔ*G* prediction methods.

The S669 dataset was reported as contaminated by considerable incorrect entries as well as mutations interfering with protein-protein or protein-ligand binding^72^. These authors composed a cleaner subset of S669, namely the S461 dataset, which retains 461 single-point mutations from the original S669 dataset with labels corrected. In this work, S461 is also used as an auxiliary benchmark test set for the performance evaluation of ΔΔ*G* prediction methods.

#### S783

S783 is a subset of S921, which was first introduced by Chen et al.^49^ and has long been a recognized test dataset for ΔΔ*G* prediction models trained on S2648. Performance of GeoDDG is also tested on this dataset to fairly compare with the existing models built purely from S2648. We filtered redundant entries from S921 and got S783 as the test set for several reasons. First, there are completely identical entries between S921 and S2648, which need exclusion. Second, some entries in S921 are simply the reverse mutations of the same protein entries in S2648. These data and their corresponding partners in the training set are thus identical for GeoDDG.

#### S8754

The S8754 dataset, composed of 8,754 single-point mutations of 301 proteins, is generated in this work as the training set for the ΔΔ*G* prediction. We collected data from two ΔΔ*G* databases, ProThermDB and ThermoMutDB, cautiously cleaned each individual piece of data, combined non-overlapping parts, and then manually verified the overlapping parts to guarantee the uniqueness and high level of credibility for each entry. Notably, we replenished the critical experimental conditions including pH and temperature for each entry of ΔΔ*G* data. After removing data redundant with the test set (*i.e*. sequence identity > 25% with the S669 dataset), we finally constructed the largest ΔΔ*G* dataset at present for model training.

#### M1261

The M1261 dataset, composed of 792 pieces of double mutation data and 469 pieces of triple or higher-order mutation data from 133 proteins, is generated in this work to evaluate the model performance in the prediction of ΔΔ*G* values of multiple mutations. Since multiple mutation data are completely absent from the training set of GeoDDG models, benchmark on this dataset could be used to evaluate the zero-shot prediction behavior and model generalizability.

#### S1626

The S1626 dataset^5^ is a Δ*T*_*m*_ dataset consisting of 1,626 single-point mutations of 95 proteins collected from the ProTherm database. It has been widely employed as the training set in prior softwares for Δ*T*_*m*_ prediction and is considered the most extensively employed dataset of its kind.

#### S571

The S571 dataset, composed of 571 single-point mutations of 37 proteins, is generated in this work as the benchmark test set for the evaluation of Δ*T*_*m*_ prediction methods. We collected data from two Δ*T*_*m*_ databases, ProThermDB and ThermoMutDB, and performed data cleaning in an approach similar to S8754. For each entry, we also replenished experimental pH information. After eliminating data redundant with the S1626 dataset (*i.e*. sequence identity > 25%), this dataset finally becomes a benchmark test set for the fair evaluation of Δ*T*_*m*_ prediction methods, since most prior Δ*T*_*m*_ prediction softwares utilized S1626 as the training set.

The S4346 dataset, composed of 4346 single-point mutations of 349 proteins, is generated in this work as the training set for the Δ*T*_*m*_ prediction. The data were collected and checked just as in the construction of S571. After removing data redundant with the test set (*i.e*. sequence identity > 25% with the S571 dataset), we finally constructed the largest Δ*T*_*m*_ dataset at present for model training.

### Information processing in the geometric-learning-based model architecture

As shown in Figure 2a, the Geometric Encoder adopts the GAT architecture, using the nodes and edges to process input features from the sequence and structure of a protein, respectively. The sequence features are derived from the large-scale protein language model ESM-2^30^, which has been improved in terms of model architecture, training parameters and training data compared to its predecessor, ESM-1b^29^, in order to capture global context information. Hence, each node in the GAT is initialized mainly by the embedding of the corresponding residue derived from the ESM-2 output. Following the idea from a prior work^73^, in our geometric learning model, each edge is initialized by the relative geometric relationship between a pair of residues derived from the protein 3D structure. The implementation details for the calculation of relative coordinates and geometries could be found in **Supplementary Methods**.

After initialized by sequence and structural features, the node embedding **x**_*i*_ and the edge embedding **z**_*ij*_ are further fused in each layer of the Geometric Encoder by

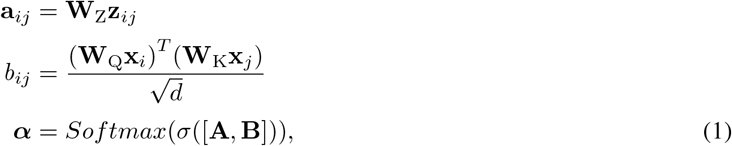

Where:

- *d* is the dimension of embedding per head in the multi-head attention mechanism,
- **A** and **B** denote the matrices composed of **a**_*ij*_ and *b*_*ij*_, respectively,
- **W**_Z_, **W**_K_ and **W**_Q_ denote three different linear transformations,
- [• ,•] denotes the concatenation operation,
- *σ* is a transformation of *σ*(**X**) = *LeakyReLU* (*Conv*2*d*(*InstanceNorm*(**X**))),
- ***α*** is the computed attention matrix.

Finally, the node embedding **x**_*i*_ and the edge embedding **z**_*ij*_ are updated through neighboring layers by

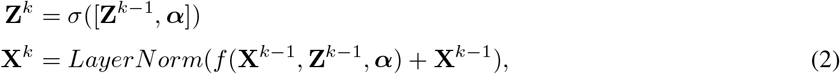

Where:

- *k* denotes the *k*^*th*^ layer of the Geometric Encoder,
- **X** and **Z** stand for the vector and matrix composed of **x**_*i*_ and **z**_*ij*_, respectively,
- *f* is an abstract function referring to a series of operations for the self-attention mechanism (see **Supplementary Methods** for details).

**X** generated from the last layer represents the 1D feature extracted by the Geometric Encoder, which can be used for downstream predictions, *e.g*., by the MLP module in GeoFitness (Figure 2b).

In the model architecture of GeoDDG and GeoDTm (Figure 2c), wide-type features and mutant features are processed by the same neural network with shared weights. Hence, difference between the output values of the wide-type and mutant proteins is supposed to change sign when their input features are switched, a design that naturally guarantees the permutational anti-symmetry of prediction results.

### Training details

We used the DMS database to train, validate and test GeoFitness. The concrete division of DMS data could be found in the **database and dataset** section. Effects of data splitting strategies for the training of GeoFitness model could be found in Figure S2 and **Supplementary Methods**. We utilized S8754 as the training set of GeoDDG and S669 as the test set for ΔΔ*G* prediction. Similarly, we utilized S4346 as the training set of GeoDTm and S571 as the test set for Δ*T*_*m*_ prediction. Notably, S8754 and S4346 are the enriched training sets collected in this work, and S571 constructed in this work is the first benchmark test set for the fair evaluation on Δ*T*_*m*_ prediction. As mentioned previously, redundancy between training sets and test sets have been carefully removed, using a stringent criterion: no proteins in the training set have sequence identity > 25% to those in the test set.

Thanks to the success of AlphaFold2 in the field of protein structure prediction, it is possible to generate accurately predicted wild-type structures from protein sequences. It was reported that models utilizing structures predicted by AlphaFold2 (pLDDT > 90) and real experimental structures could achieve similar prediction accuracies in the prediction of mutation effects^74^. Nevertheless, we trained two versions of GeoDDG/GeoDTm, utilizing experimental wild-type structures (GeoDDG-3D/GeoDTm-3D) and AlphaFold2-predicted wild-type structures (GeoDDG-Seq/GeoDTm-Seq) for training, respectively. The details of input features are briefly stated below, summarized in Table S2 and detailedly described in **Supplementary Methods**.

We selected the 33-layer 650M-parameter model of ESM-2^30^ to generate the “word” embedding for each residue. What’s more, FoldX^41^ software was utilized to predict the structures of mutants based on the wild-type structure. We only selected the 32 residues closest to the mutation location in the Cartesian space as the considered residues for structural feature extraction.

In the optimization of GeoFitness, in order to utilize the multi-labeled DMS data for model training, we converted all fitness data into the relative ranks for all possible mutations at each individual position, regardless of the original meaning of each fitness indicator. This pre-processing precedure unifies labels in the DMS database and simultaneously retains the key information (*i.e*. the relative ranking between mutations) required in practical protein engineering. We took the Spearman correlation coefficient, a metric relying on ranks, as the loss function for model training. Particularly, we adopted the soft Spearman correlation coefficient developed by Blondel *et al*., which unlike the traditional metric, is differentiable and thus allows the gradient update in the neural network^75^ (see **Supplementary Methods** for a brief introduction). This strategy enables the training of a unified GeoFitness model utilizing multi-labeled DMS data in this work. In the model training of GeoDDG and GeoDTm, parameters of the pre-trained geometric encoder inherited from GeoFitness were initially frozen to allow the rapid optimization of the other parameters. Subsequently, all parameters were allowed to change slightly in the fine-tuning stages. Furthermore, we adopted a conjugated loss function here for the ΔΔ*G* and Δ*T*_*m*_ predictions, which includes both the soft Spearman correlation coefficient and the mean square error (MSE) between predicted and experimental values. The detailed training procedure is described in **Supplementary Methods**.

### Evaluation metrics

Pearson correlation coefficient (indicated by *r*) is defined as

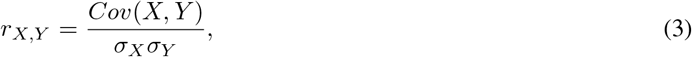

where *Cov* represents the covariance and *σ* stands for the standard deviation.

Spearman correlation coefficient (indicated by *ρ*) is calculated similarly to Pearson correlation coefficient *r* but is applied on the rank

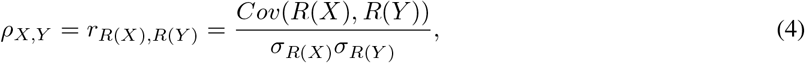

where *R* denotes the rank of a variable.

In this work, we mainly used the Spearman correlation coefficient, Pearson correlation coefficient, root mean square error (RMSE) and mean absolute error (MAE) between the predicted and experimental ΔΔ*G*/Δ*T*_*m*_ values to evaluate the performance of the methods. Additionally, we adopted two evaluation indicators to assess the ability of the predictors to reproduce the anti-symmetric property of ΔΔ*G* and Δ*T*_*m*_, namely *r*_*d*−*i*_ and ⟨*δ*⟩, defined as

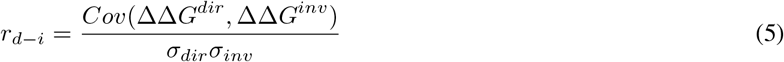

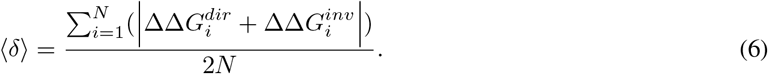

*r*_*d*−*i*_ is the Pearson correlation coefficient between the direct (*i.e*. wild-type-to-mutant) and the corresponding inverse (*i.e*. mutant-to-wild-type) sequence variations, while ⟨*δ*⟩ quantifies the bias between direct and inverse prediction values. A method that perfectly conforms to anti-symmetry should have *r*_*d*−*i*_ equal to -1 and ⟨*δ*⟩ equal to 0.

## Results

### Evaluation of protein fitness prediction by GeoFitness

As mentioned in **Methods**, we unified the multi-labeled fitness data into the relative rankings of available mutations at each individual position for each protein to facilitate the model training of GeoFitness. Therefore, the final GeoFitness is a universal fitness prediction model, which does not need to be retrained using extra fitness data of the target protein before practical prediction, *i.e*. allowing zero-shot prediction.

Here, we evaluated the performance of GeoFitness using the DMS database (DeepSequence + MaveDB). Specifically, for each protein in the DMS database, we predicted the fitness scores of all candidate single mutations in the test set, calculated the Spearman correlation coefficient between predicted and experimental values at each individual position, and averaged this value for the whole protein.

We first compared GeoFitness with other unified models that are completely independent of fitness data, including RF_joint_, MSA Transformer, ESM-1b, ESM-1v and ESM-2. As shown in Figure 3, GeoFitness remarkably outperforms these unsupervised models, approaching higher Spearman correlation coefficient for almost all proteins in the DMS database, which indicates that the use of fitness data in model training significantly improves the model performance. Subsequently, we continued the evaluation against ECNet and SESNet, two state-of-the-art methods that need to train a specific model for each individual protein. Despite the unfairness of such a comparison to our method, GeoFitness exhibits better performance on more than half of the proteins tested (Figure 3 and Figure 4a). When averaged over all proteins in the DMS database, the universal GeoFitness model reaches a slightly higher Spearman correlation coefficient than the series of specifically optimized ECNet and SESNet models (0.62 vs. 0.59 and 0.57). Interestingly, the advantage of GeoFitness over ECNet enlarges significantly with the reduction in the training data scale (Figure S3). Particularly, GeoFitness exhibits relatively robust performance in the few-shot prediction scenario when less than 10% of the data are used for model training and preserves an acceptable prediction power even in the zero-shot prediction scenario (Figures S2b and S3, see the discussion in **Supplementary Model Evaluation** for details). Therefore, although information is lost by the transformation of absolute fitness data into relative rankings in the optimization of GeoFitness, the large-scale enrichment of multi-labeled training data not only compensate this negative impact but also remarkably improve the model generalizability.

**Figure 3.**
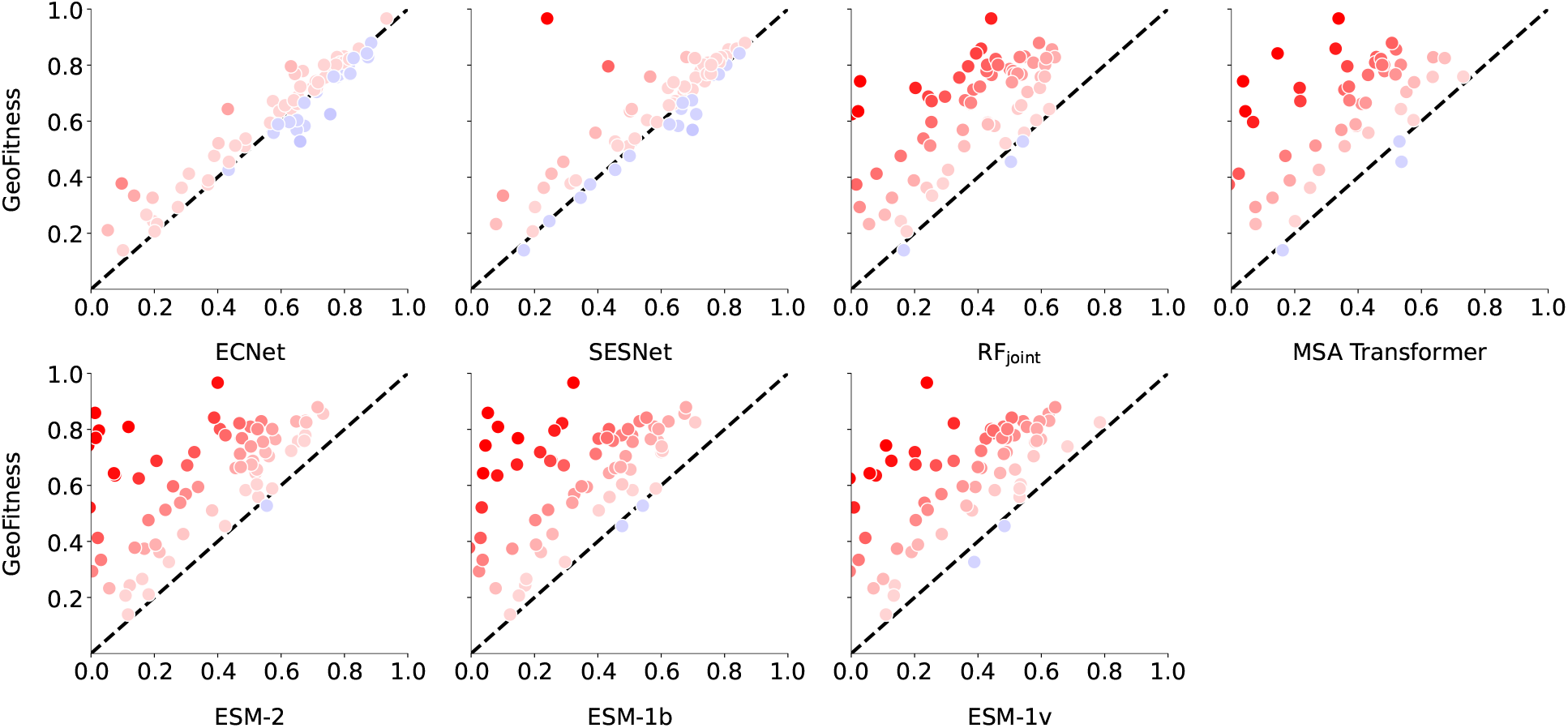
Pairwise comparisons of GeoFitness with other methods for the prediction of mutation effects on protein fitness. Each point in the figure represents the evaluation results of one protein in the DMS database using the metric of Spearman correlation coefficient. The point is colored red if GeoFitness outperforms the other method and blue otherwise, with the degree of opaqueness reflecting the magnitude of difference. Notably, metrics in this figure are computed on test subsets of DMS data rather than the full DMS dataset.

**Figure 4.**
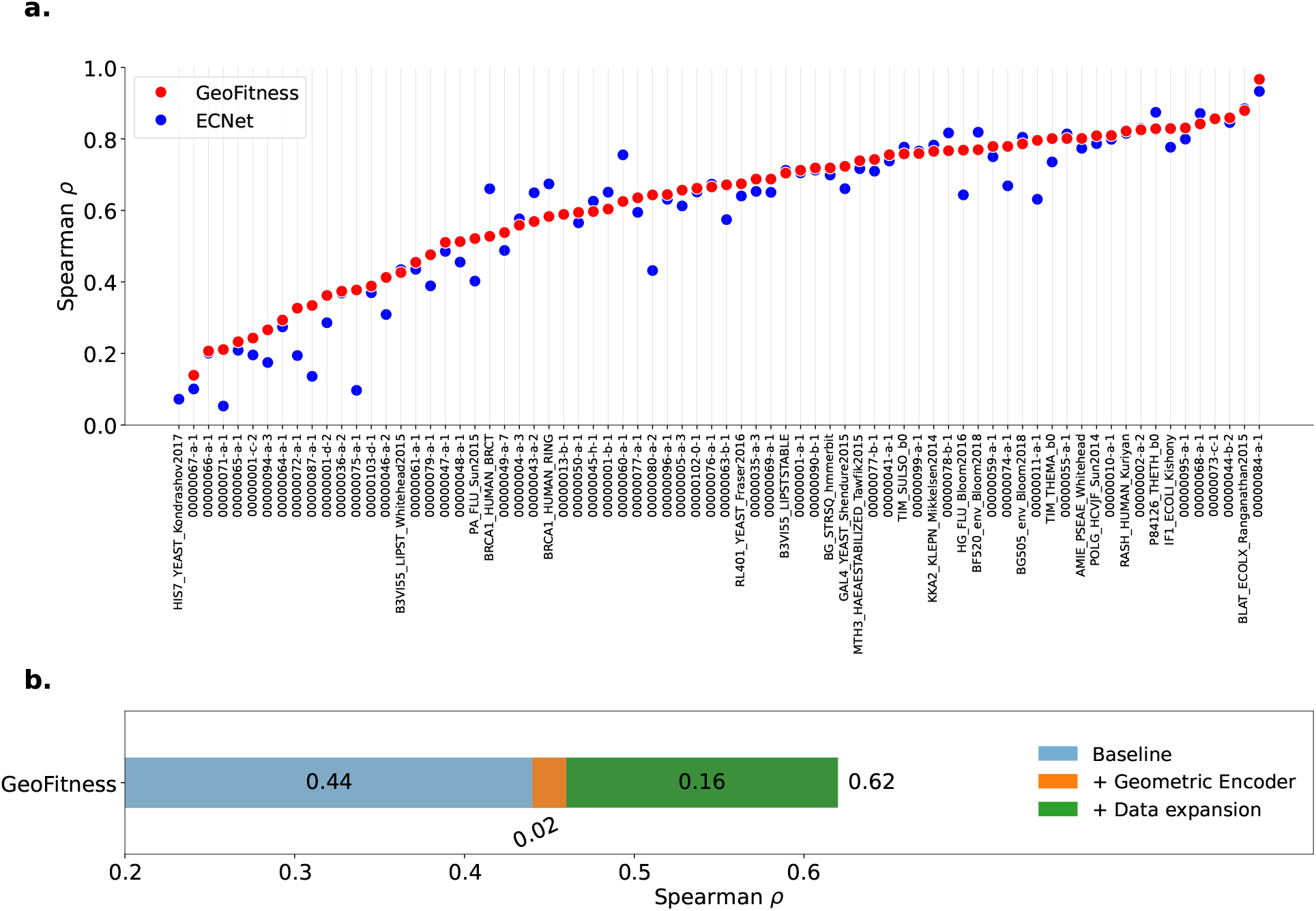
Detailed analysis for GeoFitness. **a)** Comparison between GeoFitness and ECNet on the DMS database. GeoFitness has higher Spearman correlation coefficient on 60 out of the 74 proteins. **b)** Ablation study of GeoFitness. Notably, metrics in this figure are computed on test subsets of DMS data rather than the full DMS dataset.

Furthermore, we conducted an ablation study of GeoFitness to analyze the contributions of model architecture and data enrichment, where the baseline model was trained using MLP as the network structure and DeepSequence as the training set. As shown in Figure 4b, the geometric-learning-based architecture slightly enhances the performance, whereas enrichment of data from the MaveDB database makes a more significant contribution, increasing the mean Spearman correlation coefficient by 0.16.

Although GeoFitness is originally designed for the prediction of single mutational effects only, we propose a procedure to make use of this model for the inference of multiple mutational effects (Figure S4). According to the preliminary testing results on double mutations collected from 7 proteins in the DMS database, the purely inferred fitness scores show considerable correlation with experimental values: the mean Spearman correlation coefficient is 0.52 and 0.48 when averaged over variants and over proteins, respectively (Table S3, details in **Supplementary Model Evaluation**).

In summary, through effective utilization of the large scale of multi-labeled fitness data, GeoFitness successfully learns the mutational effects without sacrificing model usability. Unlike the fitness prediction, the amount of labeled data is typically reduced by at least one order of magnitude in downstream tasks like the protein stability prediction. Considering the biological relevance between protein fitness and stability, we reused the pre-trained geometric encoder of GeoFitness as information extractor (with inherited network architecture and parameter values) in the GeoDDG/GeoDTm models for ΔΔ*G*/Δ*T*_*m*_ predictions (Figure 2b and Figure 2c). Notably, the lack of label overlap between the DMS fitness data and protein stability data safely eliminates the possibility of information leaking by this reutilization.

### Evaluation of ΔΔ*G* prediction by GeoDDG

GeoDDG uses a module with shared weights to process the sequence and structural information of both wild-type and mutant proteins and takes their difference for prediction. Unlike GeoFitness that is mainly applicable for single mutation effects, GeoDDG is, in principle, able to predict the change of Δ*G* caused by an arbitrary number of mutations, although the model in this work is trained by single mutation data only.

Available ΔΔ*G* prediction methods can be classified into sequence-based and structure-based, depending on whether the experimental structural information is required for inference. Here, we trained two versions of GeoDDG, namely GeoDDG-Seq and GeoDDG-3D, which retrieve information from AlphaFold2-predicted structures and experimental structures, respectively, and evaluated their performance against mainstream state-of-the-art methods on the S669 benchmark test set. Notably, in this work, redundancies in training sets of GeoDDG have been strictly removed against the test set S669 to guarantee fair evaluation (see **Methods** for details), and more importantly, the zero overlap between the training set of the pre-trained GeoFitness model and S669 (Figure S1) precludes the possibility of information leaking during performance evaluation.

As shown in Table 1, GeoDDG-Seq and GeoDDG-3D significantly outperform the other methods of their respective categories in all metrics. Particularly, our methods exhibit an advantage of a large margin in terms of the Spearman correlation coefficient for the direct mutational effects, with relative improvements of 36.4% (0.60 vs. 0.44, see Figure 5) and 32.6% (0.61 vs. 0.46, see Figure 6) over the second best methods in the sequence-based and structure-based categories, respectively. Moreover, GeoDDG is the only method in this test that can reduce the absolute prediction error (*i.e*. MAE) of ΔΔ*G* to be less than 1 kcal/mol. Additionally, both versions of GeoDDG successfully ensure the anti-symmetric ΔΔ*G* values between the direct and inverse mutations, as demonstrated by *r*_*d*−*i*_ and ⟨*δ*⟩ values of -1 and 0, respectively. More detailed discussion about the preservation and break of the anti-symmetry as well as additional tests on the S^sym76^ and S2000^77^ sets could be found in **Supplementary Model Evaluation**.

**Table 1.** Comparison of GeoDDG with exsiting models on the S669 dataset.

| Method | Total |  | Direct |  | Inverse |  | Antisymmetry |  |
| --- | --- | --- | --- | --- | --- | --- | --- | --- |
| | $\rho/r$ | RMSE/MAE | $\rho/r$ | RMSE/MAE | $\rho/r$ | RMSE/MAE | $r_{d-i}$ | $\langle\delta\rangle$ |
| *Sequence-based* |  |  |  |  |  |  |  |  |
| INPS-Seq | 0.64/0.61 | 1.52/1.10 | 0.44/0.43 | 1.52/1.09 | 0.44/0.43 | 1.53/1.10 | **-1.00** | 0.03 |
| ACDC-NN-Seq | 0.61/0.59 | 1.53/1.08 | 0.43/0.42 | 1.53/1.08 | 0.43/0.42 | 1.53/1.08 | **-1.00** | **0.00** |
| DDGun | 0.59/0.57 | 1.76/1.27 | 0.43/0.40 | 1.75/1.27 | 0.41/0.38 | 1.77/1.27 | -0.96 | 0.06 |
| I-Mutant3.0-Seq | 0.36/0.37 | 1.92/1.47 | 0.35/0.34 | 1.56/1.16 | 0.23/0.22 | 2.22/1.79 | -0.48 | 0.76 |
| MuPro | 0.31/0.32 | 2.03/1.58 | 0.26/0.25 | 1.61/1.21 | 0.22/0.20 | 2.38/1.96 | -0.32 | 0.95 |
| SAAFEC-Seq | 0.19/0.26 | 2.01/1.53 | 0.35/0.36 | 1.54/1.12 | -0.01/-0.01 | 2.40/1.94 | -0.03 | 0.83 |
| GeoDDG-LE-Seq | 0.68/0.67 | 1.40/1.00 | 0.54/0.53 | 1.40/1.00 | 0.54/0.53 | 1.41/1.00 | **-1.00** | **0.00** |
| GeoDDG-Seq | **0.71/0.70** | **1.37/0.97** | **0.60/0.58** | **1.37/0.97** | **0.60/0.58** | **1.37/0.97** | **-1.00** | **0.00** |
| *Structure-based* |  |  |  |  |  |  |  |  |
| DDMut | 0.63/0.62 | 1.49/1.06 | 0.44/0.44 | 1.49/1.06 | 0.44/0.44 | 1.49/1.07 | -0.90 | 0.14 |
| PremPS | 0.63/0.61 | 1.50/1.07 | 0.42/0.41 | 1.51/1.09 | 0.42/0.42 | 1.49/1.05 | -0.85 | 0.19 |
| ACDC-NN | 0.63/0.61 | 1.50/1.05 | 0.45/0.46 | 1.49/1.05 | 0.44/0.45 | 1.50/1.06 | -0.98 | 0.04 |
| INPS3D | 0.57/0.55 | 1.64/1.19 | 0.44/0.43 | 1.50/1.07 | 0.36/0.33 | 1.76/1.31 | -0.50 | 0.51 |
| DDGun3D | 0.55/0.57 | 1.61/1.13 | 0.43/0.43 | 1.60/1.11 | 0.41/0.41 | 1.62/1.14 | -0.97 | 0.09 |
| DynaMut | 0.48/0.50 | 1.65/1.21 | 0.37/0.41 | 1.60/1.19 | 0.36/0.34 | 1.69/1.24 | -0.58 | 0.31 |
| ThermoNet | 0.46/0.51 | 1.64/1.20 | 0.37/0.39 | 1.62/1.17 | 0.34/0.38 | 1.66/1.23 | -0.85 | 0.16 |
| PopMusic | 0.43/0.46 | 1.82/1.36 | 0.41/0.41 | 1.51/1.09 | 0.22/0.24 | 2.09/1.64 | -0.32 | 0.71 |
| FoldX | 0.41/0.31 | 2.40/1.53 | 0.28/0.21 | 2.32/1.57 | 0.34/0.22 | 2.48/1.50 | -0.20 | 0.75 |
| MAESTRO | 0.39/0.44 | 1.80/1.35 | 0.46/0.50 | 1.44/1.06 | 0.19/0.20 | 2.10/1.65 | -0.22 | 0.66 |
| DUET | 0.39/0.41 | 1.86/1.39 | 0.42/0.41 | 1.52/1.10 | 0.23/0.23 | 2.14/1.68 | -0.12 | 0.74 |
| mCSM | 0.34/0.37 | 1.96/1.49 | 0.36/0.36 | 1.54/1.13 | 0.21/0.22 | 2.30/1.86 | -0.05 | 0.87 |
| I-Mutant3.0 | 0.28/0.32 | 1.96/1.49 | 0.35/0.36 | 1.54/1.12 | 0.15/0.15 | 2.32/1.87 | -0.06 | 0.81 |
| SDM | 0.28/0.32 | 1.93/1.45 | 0.39/0.41 | 1.67/1.26 | 0.14/0.13 | 2.16/1.64 | -0.40 | 0.48 |
| GeoDDG-LE-3D | 0.70/0.68 | 1.39/0.98 | 0.57/0.55 | 1.39/0.98 | 0.57/0.55 | 1.39/0.98 | **-1.00** | **0.00** |
| GeoDDG-3D | **0.73/0.71** | **1.33/0.96** | **0.61/0.60** | **1.33/0.96** | **0.61/0.60** | **1.33/0.96** | **-1.00** | **0.00** |

The winner in each metric is highlighted in bold. "Direct" refers to the mutations from the wild type to mutant, "Inverse" means the corresponding mutations in the reverse direction, and "Total" stands for the combination of the above categories. Here, $\rho$ and $r$ denote Spearman and Pearson correlation coefficients, respectively. Details of all metrics are described in **Methods**. The raw results of the other models except DDMut<sup>50</sup> are taken from a prior work<sup>52</sup>.

**Figure 5.**
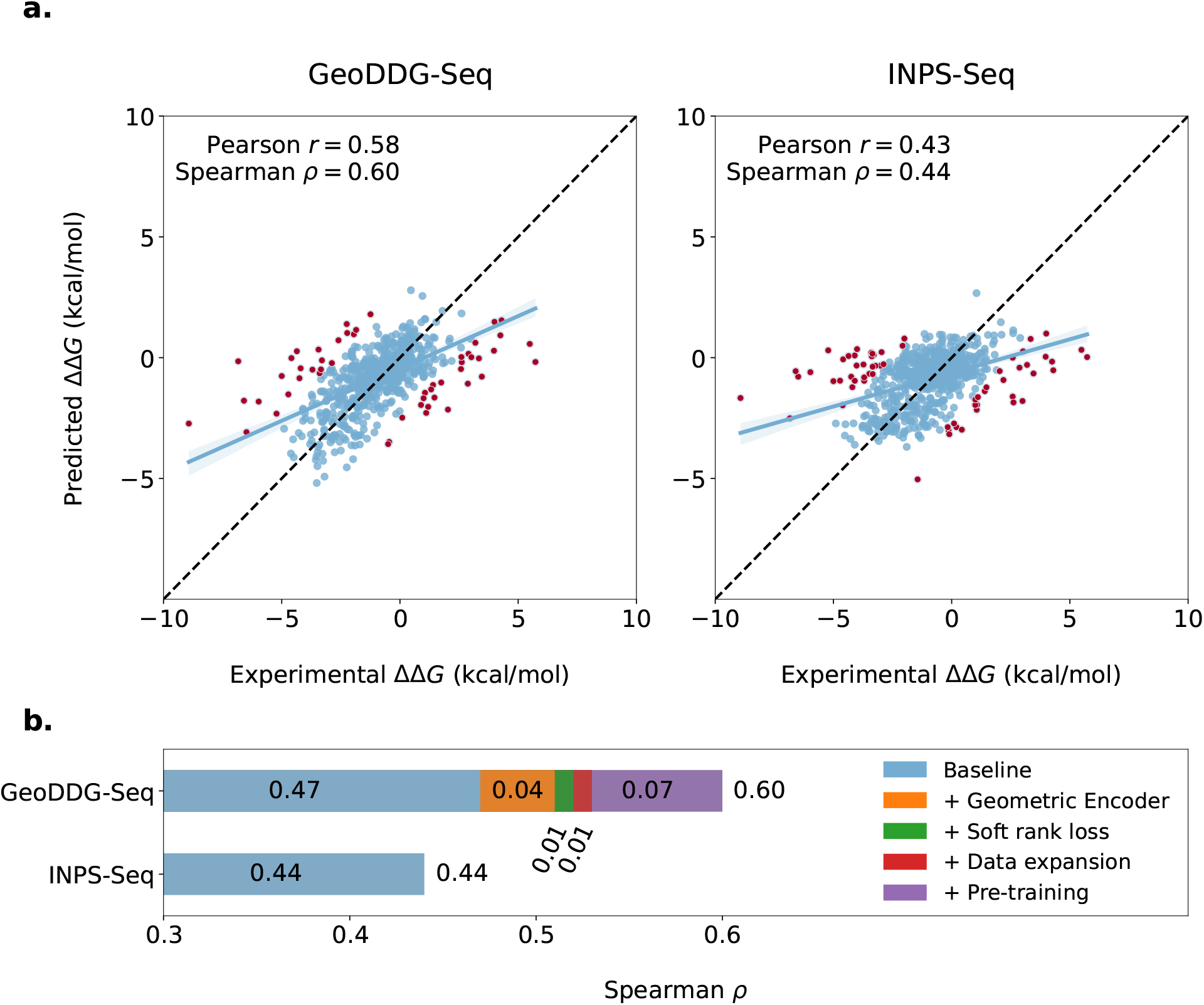
Detailed analysis for GeoDDG-Seq. **a)** Side-by-side comparison between GeoDDG-Seq and the second best sequence-based predictor, INPS-Seq, on S669. Predictions with error > 2.5 kcal/mol are identified as outliers (49 in GeoDDG-Seq vs. 54 in INPS-Seq) and are colored red in the figures. **b)** Ablation study of GeoDDG-Seq.

**Figure 6.**
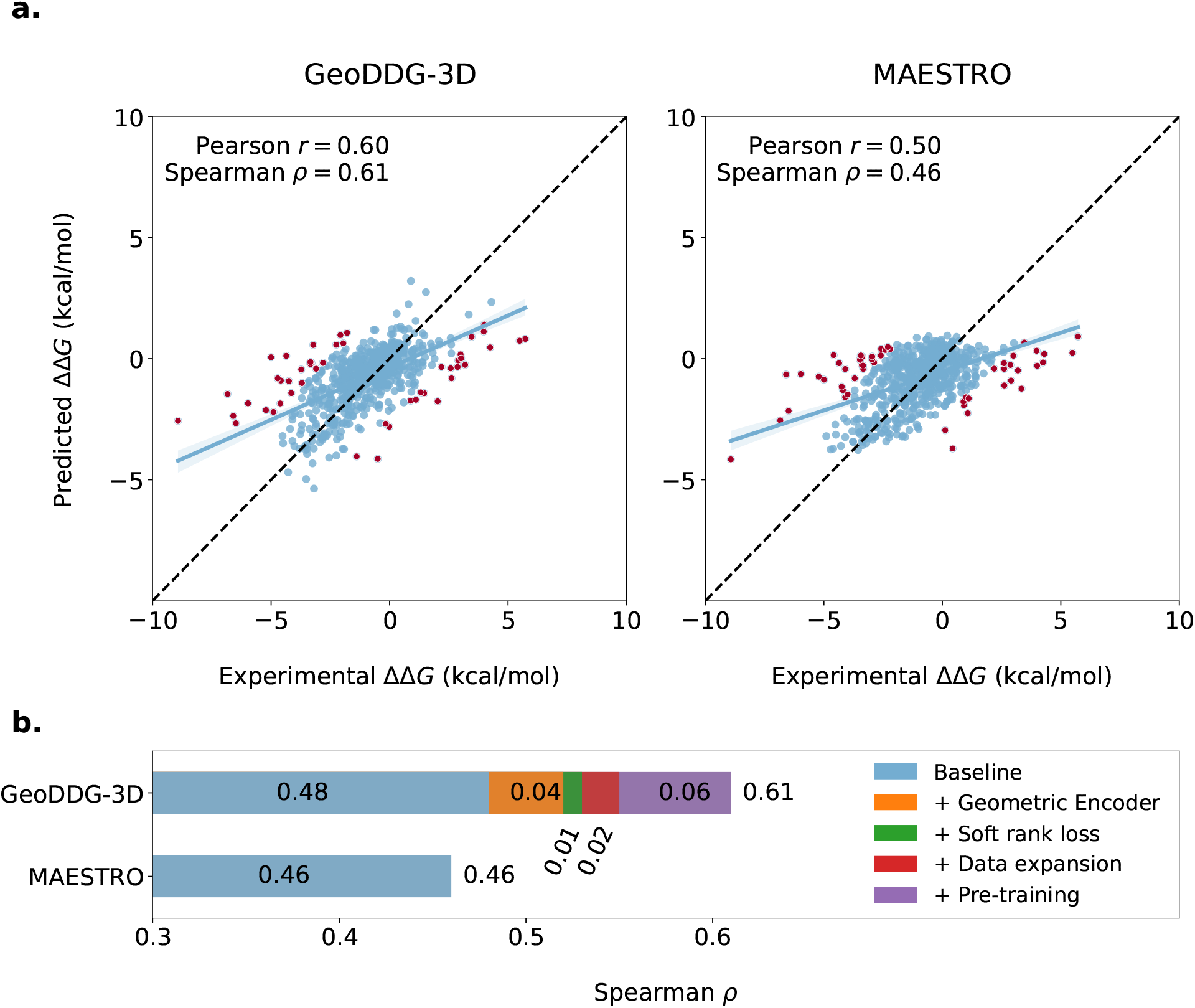
Detailed analysis for GeoDDG-3D. **a)** Side-by-side comparison between GeoDDG-3D and the second best structure-based predictor, MAESTRO, on S669. Predictions with error > 2.5 kcal/mol are identified as outliers (52 in GeoDDG-3D vs. 64 in MAESTRO) and are colored red in the figures. **b)** Ablation study of GeoDDG-3D.

In a recent work^72^, the authors claimed that the S669 dataset was contaminated and composed a cleaner subset, namely S461 (see **Methods** for details). Hence, we also conducted evaluation in the S461 dataset. As expected, nearly all methods exhibit enhanced performance in this subset (Table S4). However, GeoDDG still shows an marked advantage over all of the other methods in nearly all metrics. Specifically, GeoDDG-Seq and GeoDDG-3D outperform the corresponding second best methods by large margins in terms of the Spearman correlation coefficient for the direct mutational effects: 0.68 vs. 0.55 and 0.67 vs. 0.60.

To further validate the importance of the innovation points of our GeoDDG model, principally the pre-training strategy and the design of network architecture, we retrained GeoDDG using limited training data, *i.e*. on the basic S2648 dataset, from which many of the previous methods were developed, and named this model as GeoDDG-LE. As shown in Table 1, when evaluated on the S669 test set, the model performance of GeoDDG-LE declines slightly in comparison with the full version GeoDDG, but still prevails all of the other methods by a large margin (*e.g*., at least 0.1 in the Spearman correlation coefficient for direct mutations), for both sequence-based and structure-based predictors. We also tested GeoDDG-LE against previous methods developed purely from the basic S2648 training set on the S783 dataset (see **Methods** for details) and found the significant advantage of GeoDDG-LE models in terms of the Spearman correlation coefficient (Table S5). Results of the above tests and the consistent behaviors of GeoDDG-LE models in 5-fold and 10-fold cross validations within the S2648 training set (Figure S5) jointly indicates that our method is capable of generating reliable prediction models for the mutation effects on the protein stability, even in the situation of limited training data.

Subsequently, we conducted ablation studies for the two versions of GeoDDG in the S669 dataset. As expected, the baseline models that were trained using the architecture of MLP on the smaller S2648 dataset show similar performances to INPS-Seq (Figure 5) and MAESTRO (Figure 6), the second best methods in the sequenced-based and structure-based categories, respectively. In both cases, the introduction of geometric encoder brings an increase of 0.04 to the Spearman correlation coefficient, while the engage of soft rank loss and data expansion further enhance the performance slightly. Interestingly, the pre-training strategy introduces the largest improvement (0.07 in GeoDDG-Seq and 0.06 in GeoDDG-3D), supporting that model pre-training using the large amount of protein fitness data can effectively overcome the challenge of limited data that have perplexed the ΔΔ*G* prediction in the past years.

Recently, a new set of DMS data was released, whose data label is non-equivalent to ΔΔ*G* but related with protein stability^78^. We thus evaluated our GeoDDG model on this dataset. Results suggest that GeoDDG predictions correlate fairly well with the DMS experimental values, with the mean Spearman correlation coefficient approaching 0.63 (Figure S6, see details in **Supplementary Model Evaluation**). On the other hand, considering that GeoDDG was trained without multiple mutational data, we subsequently tested its behavior in the zero-shot prediction of the ΔΔ*G* caused by multiple mutations. Here, we composed a testing set M1261 (see **Methods**) and evaluated GeoDDG against two mainstream methods, DynaMut2^79^ and DDMut^50^, which employed data of both single and multiple mutations for model training. In brief, GeoDDG shows a comparable performance to DDMut, whereas both methods significantly prevail DynaMut2 (Table S7, see details in **Supplementary Model Evaluation**). The excellent performance of GeoDDG in the above two zero-shot prediction tasks sufficiently corroborates the good generalizability of our model.

In conclusion, based on the comprehensive evaluations described above, GeoDDG enormously improves the prediction of mutation effects on the Δ*G* of proteins through the effective use of data by the pre-training strategy and the adequate processing of protein structural information by geometric learning.

### Evaluation of Δ*T*_*m*_ prediction by GeoDTm

Similar to GeoDDG, GeoDTm can predict the influence on *T*_*m*_ of a protein by multiple mutations, with “Seq” and “3D” versions denoting the models trained using AlphaFold2-predicted structures and experimental structures, respectively.

However, unlike the ΔΔ*G* prediction, there is no previously established benchmark test set for the Δ*T*_*m*_ prediction, thus disallowing the fair evaluation of all methods. To address this problem, we curated a test set of 571 samples for Δ*T*_*m*_, named S571, which is non-redundant with currently available training datasets (see **Methods** for details) and has zero overlap with the training set of the pre-trained GeoFitness model at the same time (Figure S1). Subsequently, we evaluated the performance of GeoDTm-Seq and GeoDTm-3D on the S571 set against two available Δ*T*_*m*_ predictors, AUTO-MUTE and HoTMuSiC, which are, unfortunately, both structured-based models and unable to provide the prediction for the inverse mutations. As shown in Table 2, GeoDTm-3D prevails the other structure-based methods in all metrics. Notably, our method tremendously enhances the Spearman correlation coefficient from 0.30 to 0.51, with a relative improvement of more than 70%. Additionally, like the case in GeoDDG, both versions of GeoDTm perfectly conform to the anti-symmetric requirement of prediction results.

**Table 2.** Comparison of GeoDTm with existing models on the S571 dataset.

| Method | Total |  | Direct |  | Inverse |  | Antisymmetry |  |
| --- | --- | --- | --- | --- | --- | --- | --- | --- |
| | $\rho/r$ | RMSE/MAE | $\rho/r$ | RMSE/MAE | $\rho/r$ | RMSE/MAE | $r_{d-i}$ | $\langle\delta\rangle$ |
| *Sequence-based* |  |  |  |  |  |  |  |  |
| **GeoDTm-Seq** | **0.54/0.52** | **8.11/5.55** | **0.51/0.46** | **8.11/5.55** | **0.51/0.46** | **8.11/5.55** | **-1.00** | **0.00** |
| *Structure-based* |  |  |  |  |  |  |  |  |
| HoTMuSiC |  |  | 0.30/0.33 | 8.41/5.70 |  |  |  |  |
| AUTO-MUTE |  |  | 0.25/0.29 | 8.50/5.79 |  |  |  |  |
| **GeoDTm-3D** | **0.55/0.53** | **8.03/5.31** | **0.51/0.47** | **8.03/5.31** | **0.51/0.47** | **8.03/5.31** | **-1.00** | **0.00** |

The winner in each metric is highlighted in bold. "Direct" refers to the mutations from the wild type to mutant, "Inverse" means the corresponding mutations in the reverse direction, and "Total" stands for the combination of the above categories. Here, $\rho$ and $r$ denote Spearman and Pearson correlation coefficients, respectively. Details of all metrics are described in **Methods**. Blank region means that the corresponding result is unavailable.

We also performed ablation studies for the two versions of GeoDTm .As shown in Figure 7, the baseline models that were trained using the MLP architecture on the smaller S1626 training set already prevail the other methods (HoTMuSiC here), which were likely to be over-trained considering the significantly reduced performance on the S571 set compared with the reported cross-validation values in their original papers. The geometric encoder, soft rank loss and data expansion jointly improve the Spearman correlation coefficient by 0.06 and 0.05 for GeoDTm-3D and GeDTm-Seq, respectively. In both versions of GeoDTm, the pre-training strategy makes the largest contribution, increasing the Spearman correlation coefficient by 0.09. This significant enhancement further demonstrates the role of pre-training using large amount of related data in eliminating overfitting and improving model generalizability.

**Figure 7.**
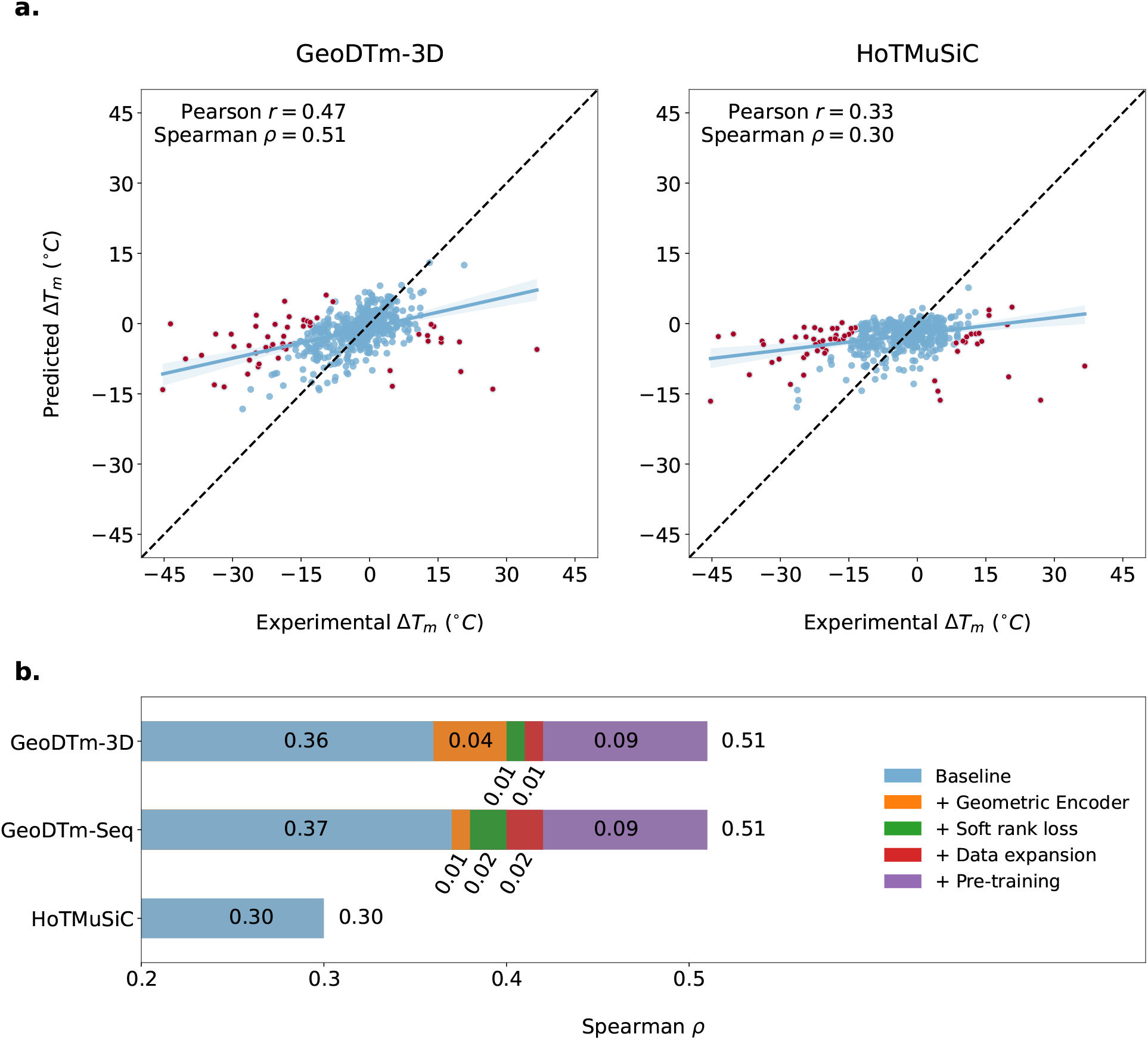
Detailed analysis for GeoDTm. **a)** Side-by-side comparison between GeoDTm-3D and the second best structure-based predictor, HotMuSiC, on S571. Predictions with error > 11.0 ^*°*^C are identified as outliers (68 in GeoDDG-3D vs. 83 in HotMuSiC) and are colored red in the figures. **b)** Ablation study of GeoDTm-3D and GeoDTm-Seq.

Hence, similar to GeoDDG, GeoDTm tremendously improves the prediction of mutation effects on the *T*_*m*_ of proteins.

## Discussion

Understanding the mutation effects is an important issue in practical protein design and engineering, and hence extensive attention has been paid to developing computational methods to learn and predict these effects accurately and efficiently. Despite the continual efforts in the model architecture design, insufficient and ineffective data utilization gradually becomes the bottleneck that prohibits the further improvement of model performance. In the fitness prediction field, although the large scale of DMS data have paved the way for the training of deep learning models, existing models have to sacrifice either the prediction power or the model usability due to the improper treatment of the multi-labeled data. In the ΔΔ*G* and Δ*T*_*m*_ prediction field, though the problem definition is clear and the data are single-labeled, the limited size and significant bias of data prohibit the training of deep learning models without overfitting. A systematic review^53^ even claimed that the average accuracy of ΔΔ*G* prediction models had not improved in the past 15 years given the evaluation results on the S669 test set^52^. The situation is even worse for the Δ*T*_*m*_ prediction, since objective test sets (like S669 in ΔΔ*G* prediction) are absent, which disallows fair evaluation for the Δ*T*_*m*_ prediction methods.

In this work, we adopted the differentiable ranking algorithm^75^ to construct a soft Spearman correlation coefficient loss, which enables the training of a unified neural network model with all DMS data regardless of their multi-labeled nature. This novel loss function allows us to make full use of the DMS dataset in the supervised training of GeoFitness, which could predict the fitness landscape of the interested protein with an accuracy higher than the state-of-the-art models without the strict requirement of model retraining using sufficient data of the same kind before practical use.

Since protein stability is closely related with protein fitness in biology, we utilized the geometric encoder of GeoFitness that was pre-trained on the fitness data to guide the prediction of ΔΔ*G* and Δ*T*_*m*_. As suggested by the ablation studies, the pre-training strategy enhances the Spearman correlation coefficient by 0.07/0.06 and 0.09 for ΔΔ*G* and Δ*T*_*m*_ prediction, respectively, indicating its marked role in alleviating the negative impact of limited data and in improving the model generalizability. Such remarkable improvements mainly arise from the correlation between the prediction targets, fitness and stability here, as well as the overwhelming data scale of the former over the latter. Moreover, the improvement introduced by the pre-training strategy is more significant in the Δ*T*_*m*_ prediction task that has less available training data than the ΔΔ*G* prediction. This observation implies the possible aids of our pre-training strategy in the methodology development of other downstream tasks such as predicting the luminescence intensity and expression level of proteins upon mutations, application tasks that are of great importance but have even less available experimental data.

To further improve the prediction powers on protein stability, we collected, cleaned and manually checked the data from ProThermDB^69^ and ThermoMutDB^70^ and curated the largest open-source protein stability datasets: S8754 for ΔΔ*G* and S4346 for Δ*T*_*m*_. Ablation studies on GeoDDG and GeoDTm support the positive impact of data expansion on model performance. Before the introduction of S669, in the ΔΔ*G* prediction field, cross validation is usually used to demonstrate the effectiveness of the model, a protocol that frequently causes bias^53^ and hinders the fair evaluation of the model performance. To address the same challenge confronted in the Δ*T*_*m*_ prediction field, we constructed a new benchmark test set S571 in this work to enable the fair model evaluation on Δ*T*_*m*_ prediction. The three newly collected datasets, S8754, S4346 and S571, are freely downloadable and may further benefit the field of protein stability prediction.

The models proposed in this work, GeoFitness, GeoDDG and GeoDTm, are all based on the geometric encoder, a geometric-learning-based network architecture that are designed to simultaneously process the information of protein sequences and structures and to enable the anti-symmetric prediction when integrated with downstream networks.

Based on the evaluation of prior models (Table 1 and Table 2), utilization of structural information clearly benefits the stability prediction (see the better performance of structure-based models vs. sequence-based ones in general), but introduces additional limitations, since experimental structures are frequently unavailable for the concerned protein target. Therefore, we trained two versions of protein stability predictors, GeoDDG/GeoDTm-3D and GeoDDG/GeoDTm-Seq, which retrieve structural information from the experimental structures and AlphaFold2-predicted ones^60^, respectively. In practical predictions, the “3D” version is suggested for better performance when the reliable experimental structure is available, while the “Seq” version is an alternative when only sequence information is known. On the other hand, whether AlphaFold2 prediction benefits the downstream protein functional prediction tasks is contraversial^74,80–84^. As shown in Figure S7, correlation between the predicted ΔpLDDT by AlphaFold2 and ΔΔ*G* is poor and is completely lost if we focus on data points of higher significance (*i.e*. absolute ΔΔ*G* value > 5 kcal/mol). Hence, the naive use of AlphaFold2 indicators like the pLDDT is unlikely to improve the functional prediction, while metrics quantifying structural changes around the mutational site may provide more insightful information. Incidentally, among the stability predictors proposed in this work, our “Seq” version can reach a prediction power very close to that of the “3D” version (Table 1 and Table 2), suggesting that proper utilization of the accurately predicted structures from AlphaFold2 as spatial constraints (*e.g*., the edge embedding in this work) can effectively assist the downstream tasks like the stability prediction, which agrees with previous findings^74^.

We have established a web server for GeoStab-suite, a suite of the three predictors GeoFitness, GeoDDG and GeoDTm. We expect the GeoStab-suite to be a useful tool for researchers in the field of protein science.

## Author contribution

Y. X. and H. G. proposed the methodology and designed the experiment. Y. X. implemented the experiment. Y. X. and D. L. analyzed the results. Y. X., D. L. and H. G. wrote the manuscript. All authors agreed with the final manuscript.

## Supplementary Materials

## 1 Supplementary Methods

### 1.1 Feature generation

Most of the features utilized are summarized in Table S2, while the 2D information calculated from the protein structure is detailedly described in this section.

The position and orientation of each residue could be delicately quantified by a local coordinate system using the positions of its C_*α*_, N and C (*i.e*. carboxyl carbon) atoms. When the local coordinate system of a given residue is taken as the reference, the global coordinates of all other residues can be transformed into relative translations (as vectors) and rotations (as matrices) to this reference frame. Here, we follow the AlphaFold2^60^ definition on the residue-level local coordinate system. Thus, the bases of the local coordinate system of the *i*^*th*^ residue, **R**_*i*_, are calculated as

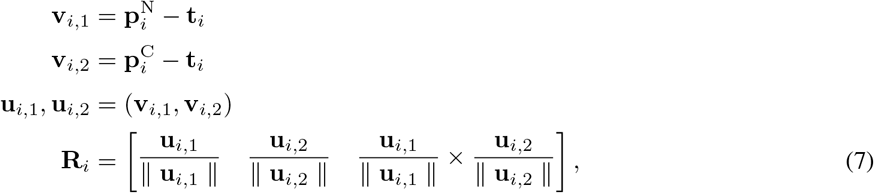

where **t**_*i*_, 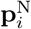 and 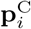 denote positions (as vectors) of the C_*α*_, N and C atoms in the global coordinate system, respectively, while the notation (·, ·) refers to the Gram-Schmidt orthogonalization operation, ∥ · ∥ stands for the metric of vector length and × denotes the cross product of vectors.

Denote the bases of the local coordinate systems of residue *a* and *b* as **R**_*a*_ and **R**_*b*_, then the rotation matrix quantifying the relative orientation of residue *b* in the local coordinate system of residue *a* can be calculated as

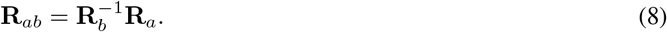

For any rotation matrix **R**, it can be converted into the quarternion form as

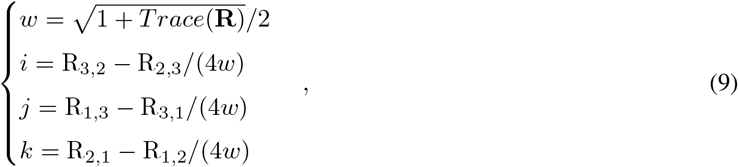

where R_*m,n*_ denotes the element in the *m*^*th*^ row and *n*^*th*^ column of the matrix **R**.

For a pair of residues *i* and *j*, we concatenate the 3-dimensional C_*α*_ relative position and the 4-dimensional relative quarternion of residue *i* in the reference coordinate system of residue *j* to compose a 7-dimensional feature so as to comprehensively describe the inter-residue relative translation and rotation. These features jointly constitute a matrix, which corresponds to the 2D information in Table S2.

### 1.2 Details for the model architecture

#### 1.2.1 Geometric encoders

##### Convolution operation in Equation 1

The geometric encoders in GeoFitness and GeoDDG/GeoDTm adopt a GAT architecture, where the node and edge embeddings are fused into the neural network following Equation 1. Specifically, the edge information arises from two sources, linear transformation of the original edge features (labeled as matrix **A**) as well as the interaction between node features through outer product (labeled as matrix **B**). These two matrices are concatenated and fed through a 2D convolution layer followed by *Softmax* to generate the attention matrix ***α***. Convolution operations are rarely seen in the traditional application of attention. Here, we briefly justify its relevance in the processing of protein information.

Traditional attentions are permutational invariant, and therefore positional encoding (PE) has to be applied to indicate relative position of a word in a sentence in natural language processing. Protein language models like ESM-1v and ESM-2 encode the positional information using periodical functions, which is justified for natural language processing but imperfect in the processing of protein sequences. Based on biophysical studies in the past years, residues pairs separated by 6 residues or fewer (named as local residues) and the other pairs (named as non-local residues) have remarkably distinct behaviors. Specifically, in a free polypeptide chain, the backbone dihedral angles of local residue pairs are correlated, whereas the correlation disappears for non-local residue pairs. In the language of biophysics, the persistence length of polypeptide chains are around 6 amino acid residues. Consequently, in many protein folding algorithm like Rosetta, local and non-local residues are frequently treated differently in the statistical potential terms. Obviously, the conventional implementation of PE by periodical functions cannot capture such a distinction between local and non-local residue pairs. In contrast, convolutional operations are enforced locally, and hence the joint application of *Conv*2*d* and PE (from ESM-2 embeddings) can fulfill the tasks of recognizing the sequence position as well as distinguishing the local and non-local residue-residue interactions simultaneously.

##### *f* function in Equation 2

Here, we provide the pseudocode for the abstract *f* function in Algorithm 1. Where:

- *L* denotes the truncation length of the proteins, chosen as 32,
- *Batch* denotes the batch size, chosen as 256,
- *node*_*dim, edge*_*dim* and *num*_*head* are hyper-parameters (see the **Hyper-parameter selection** section),
- **W** is the learnable weight matrix of the linear transformation,
- **X, Z** and ***α*** denote the input node feature, edge feature and attention matrix, respectively.

###### Algorithm 1 Definition of the abstract function *f*

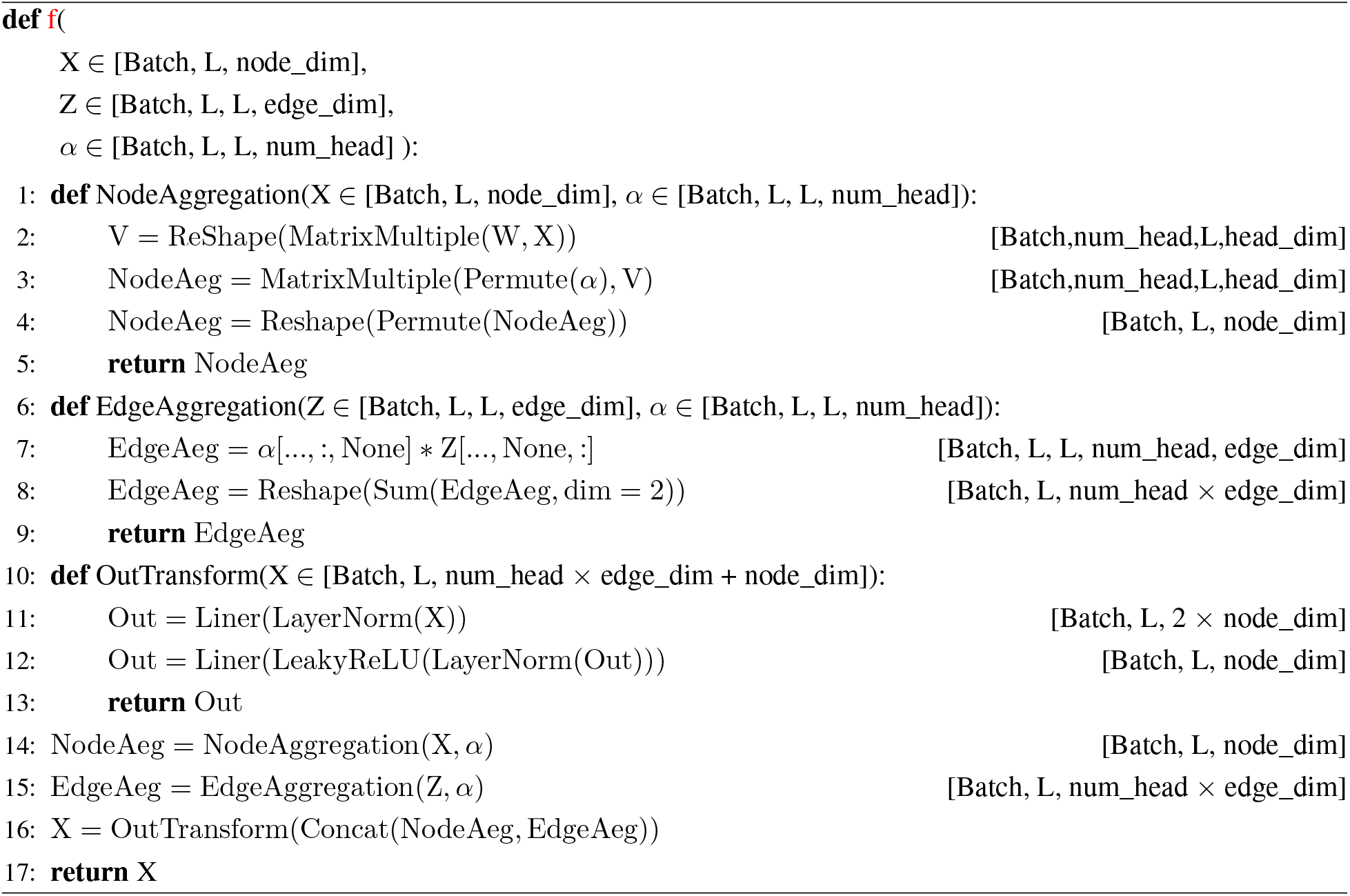

#### 1.2.2 MLP and output layers

##### GeoFitness

The MLP and output layers in GeoFitness can be formulated as

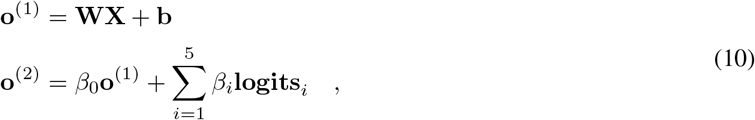

Where:

- **X** (*Batch* × *L* × *node*_*dim*) is the feature generated by the geometric encoder of GeoFitness,
- **W** is the weight matrix for the MLP layer,
- **b** is the bias vector for the MLP layer,
- **o**^(1)^ (*Batch* × *L* × 20) is the output of the MLP layer,
- *β*_*i*_ *i* ∈ {0, 1, 2, 3, 4, 5} are learnable weight parameters,
- **logits**_*i*_ *i* ∈ {1, 2, 3, 4, 5} are the logits predicted by five ESM-1v models,
- **o**^(2)^ is the final output corresponding to the *L* × 20 fitness landscape.

##### GeoDDG and GeoDTm

The MLP and output layers in GeoDDG/GeoDTm can be formulated as

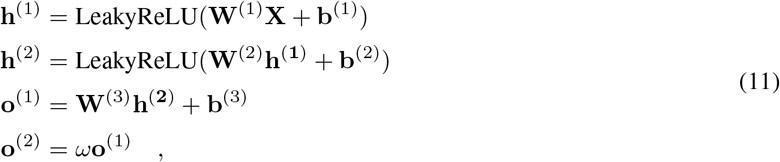

Where:

- **X** (*Batch* × *L* × *node*_*dim*) is the feature generated by the geometric encoder of GeoDDG/GeoDTm,
- **W**^(*l*)^ is the weight matrix for MLP layer *l*,
- **b**^(*l*)^ is the bias vector for MLP layer *l*,
- **h**^(1)^ (*Batch* × *L* × *node*_*dim*) is the activation output for MLP layer 1,
- **h**^(2)^ 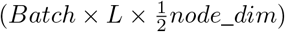 is the activation output for MLP layer 2,
- **o**^(1)^ (*Batch* × *L* × 1) is the output of the MLP layers,
- *ω* is a learnable scaling parameter, initialized as 1,
- **o**^(2)^ is the final readout for an input wild-type/mutant protein.

ΔΔ*G*/Δ*T*_*m*_ is computed from the difference between the wild-type and mutant readouts.

#### 1.2.3 Hyper-parameter selection

The hyper-parameters awaiting selection include dimension of node features (*node*_*dim*), dimension of edge features (*edge*_*dim*), number of multi-head attention matrices (*num*_*head*) and number of GAT layers in the geometric encoder (*num*_*layer*). The detailed results of the hyper-parameter selection process are exemplified for GeoDDG in Table S8, where the hyper-parameters are scanned and evaluated by grid search and the combination leading to the best performance in validation is chosen for the final model. The selection processes of GeoFitness and GeoDTm are similar. The numbers of parameters in the final models of GeoFitness, GeoDDG-3D, GeoDDG-Seq, GeoDTm-3D and GeoDTm-Seq are 1.11 million, 3.16 million, 3.88 million, 3.27 million and 2.46 million, respectively.

Besides, the hyper-parameter *ϵ* of the soft Spearman correlation coefficient is also tunable. However, we chose the default value as suggested by the original paper of the soft ranking algorithm (see the **loss function** of the GeoFitness section for details). In the training procedure of GeoDDG/GeoDTm, the sequence of parameter freezing and optimization as well as the application protocol for the conjugate loss could be regarded as additional hyper-parameters, whose description could be found in the **training details** of the GeoDDG/GeoDTm section.

In addition to the hyper-parameters, factors like the weight initialization scheme as well as the use of multiple random seeds also influence the model performance, as exemplified for GeoDDG in Table S9. Based on the results, we finally chose the default weight initialization. Considering the minor effects of random seeds (as suggested by the small standard deviation of the multiple repetitions), we abandoned the use of multiple random seeds so as to accelerate the inference process.

### 1.3 GeoFitness training procedure

#### 1.3.1 Feature utilization

The 1D information provided by ESM-2 (esm2_t33_650M_UR50D) and the 2D information calculated from protein structure are taken as the input node and edge embeddings of GeoFitness, respectively. In addition, the logits predicted by five ESM-1v models (esm1v_t33_650M_UR90S_1, esm1v_t33_650M_UR90S_2, esm1v_t33_650M_UR90S_3, esm1v_t33_650M_UR90S_4 and esm1v_t33_650M_UR90S_5) are taken as the weights for point mutations at each position, which in combination with the residue-level output from the MLP layers of GeoFitness produce the final fitness landscape by weighted averaging (Equation 10).

#### 1.3.2 Dataset splitting

As stated in **Methods**, the DMS data of each protein were randomly assigned as training, validation and testing following the 7:1:2 ratio. The splitting of data could be performed in two different manners, *i.e*. the randomized split and position-specific split as shown in Figure S2a. Theoretically, model training following the latter scheme is less prone to the potential information leak between the various mutations upon the same individual residue. However, when evaluated by the 10-fold cross validation, the model derived from this strategy did not achieve satisfying performance (Figure S2b). Particularly, the variation of model performance between different cross-validation processes became significantly larger than the randomized splitting strategy. A possible reason is that the supervised training of a universal model with distinct DMS labels is already a highly challenging task and that residue-wise masking of the labels (*i.e*. position-specific splitting) will further exacerbate the difficulty. Consequently, in this work, we still adopted the randomized splitting strategy for the training of GeoFitness. Besides, as shown in Figure S2b, we also tried a third data splitting strategy to test the model behavior in truely zero-shot prediction. Details could be found in **Supplementary Model Evaluation**.

#### 1.3.3 Loss function

To comprehensively utilize the multi-labeled DMS data for the training of a universal GeoFitness model, we converted all fitness data into the relative ranks so that the original meaning of each fitness indicator was unified as a kind of general information of protein mutation effect. We then employed the soft Spearman correlation coefficient developed by Blondel *et al*.^75^ as the main loss function to enforce the learning of the ranking information. Unlike the traditional metrics, this loss function, named as SCC loss, is differentiable and thus allows the gradient update in the neural network.

Here, we briefly introduce the basic idea behind the soft ranking algorithm and the calculation of the soft Spearman correlation coefficient. Traditional ranking should be evaluated in a discrete manner and is thus non-differentiable. In the soft ranking algorithm, ranking is first converted to an continuous optimization task in the permutahedran of descending indices [*n, n* − 1, …, 1]. A convex regularization is then applied to the optimization function, with a hyper-parameter *ϵ* to tune the “softness” of this regularization, which could improve the differentiability of the objective function. In this work, we employed the entropic regularization *E*(***μ***) = ⟨***μ***, log ***μ*** − 1⟩ with the default value of *ϵ*, since according to the original paper, the value of this hype-parameter is insensitive for the ranking task. With the soft ranks computed, we could evaluate the corresponding Spearman correlation coefficient instantly following Equation 4.

#### 1.3.4 Training details

We trained GeoFitness by taking the SCC loss as the only objective function. The Adam optimizer was chosen to train the model for 200 iterations. The initial learning rate was set as 10^−3^ and was declined by half if the validation loss stopped dropping for 10 consecutive iterations. We adopted an early stop strategy: the training process will be terminated if the validation loss ceases to decline for 20 consecutive iterations. Hence, training finished once either the maximum number of training iterations was reached or the early stop criterion was met.

### 1.4 GeoDDG and GeoDTm training procedure

#### 1.4.1 Feature utilization

We first present the feature utilization of GeoDDG. In GeoDDG, the information sequentially flow through the pre-trained GeoFitness module, the downstream geometric encoder, MLP layers and the output layer. The input features of pre-trained GeoFitness module are almost the same as described in **Feature utilization of GeoFitness** section, except that logits predicted by the five ESM-1v models are excluded. The outputs from the pre-trained GeoFitness module will be fed into the downstream geometric encoder as well as MLP and output layers of GeoDDG/GeoDTm, in combination with additional node and edge features.

The additional node features include 1D output of ESM-2 (*L* × 1280), physicochemical properties (*L* × 7), pH and temperature. The pH value is clipped to the interval of [0, 11], while the temperature is obtained by dividing the raw temperature in Celsius by 10 and then clipping the quotient to the interval of [0, 12]. The zero-dimensional features, *i.e*. pH and temperature, are expanded through replication to the shape of *L* × 1 before concatenation with physicochemical properties and ESM-2 output.

The additional edge features include the 2D information calculated from the protein structure, which is the experimental protein structure in GeoDDG-3D but is the structure predicted by AlphaFold2 in GeoDDG-Seq.

The feature utilization of GeoDTm is almost the same as that of GeoDDG except that the temperature feature is absent.

#### 1.4.2 Loss function

When training GeoDDG and GeoDTm, we adopted both the SCC loss and the traditional MSE loss to compose a conjugate loss function, which allows the simultaneous learning of both the relative ranking and the absolute values for the residue mutational effects on protein stability, namely ΔΔ*G* and Δ*T*_*m*_. The detailed application of the two loss functions in the training process is described in the **following** section.

#### 1.4.3 Training details

In GeoDDG/GeoDTm, the geometric encoder of the pretrained GeoFitness is re-utilized as the major feature extractor. The output of this geometric encoder are concatenated with the additional node and edge features, which are then fed into the downstream geometric encoder as well as the MLP and output layers of GeoDDG/GeoDTm for prediction.

If not mentioned otherwise, the basic training settings of GeoDDG/GeoDTm are the same as described in **Training details of GeoFitness** section. The overall training process is divided into three stages. In the first stage, we froze parameters of the pre-trained module (from GeoFitness) as well as the scaling parameter *ω* in the last linear projection layer as described in Equation 11 of **Details for the model architecture** section, while optimized the rest parameters of GeoDDG/GeoDTm model using the SCC loss until the maximum training iteration was reached or the early stop criteria was met. In the second fine-tuning stage, parameters in the pre-trained module were released, and thus all parameters except the scaling parameter *ω* were optimized with an initial learning rate of 10^−5^. After the first two stages of training, the relative ranking information had been fully captured by the model. To further enhance the model’s capability of predicting the absolute values of ΔΔ*G* and Δ*T*_*m*_, we utilized the MSE loss to tune the model in the third stage. Specifically, we optimized the scaling parameter *ω* using the MSE loss but kept all of the other parameters fixed. Fine-tuning of this scaling parameter in the last output layer ensured that the model could improve the prediction on the absolute value without perturbing the relative ordering across all outcomes.

Notably, in the last output layer of GeoDDG/GeoDTm, the network readout is only tuned by a scaling factor without the bias term, since the final prediction result comes from the difference between the readouts of wild-type and mutant proteins, a procedure in which fixed bias will be automatically canceled out.

## 2 Supplementary Model Evaluation

### 2.1 Generalizability of GeoFitness

In the benchmark evaluation of protein fitness prediction, both GeoFitness and ECNet were trained using 70% of the DMS data. To further verify the model generalizability, we gradually removed data from the training set, re-trained both GeoFitness and ECNet repeatedly, and then re-evaluated their performances in the original test set. As shown in Figure S3, although both models exhibit weakened prediction powers with the reduction in the training data scale, GeoFitness is significantly more robust than ECNet. Particularly, when the proportion of training data drops below 10%, a scenario corresponding to few-shot prediction, or even in the extreme case of 1% of training data, a scenario close to zero-shot prediction, GeoFitness still preserves an acceptable performance, with the mean Spearman correlation coefficient nearly approaching 0.4. The sharp contrast between GeoFitness and ECNet in the region of limited training data (*e.g*., data proportion < 10%) implies that our scheme of training a universal model using multi-labeled DMS data indeed overcomes the challenge of insufficient training data and thus effectively improve the model generalizability.

We also evaluated the prediction power of GeoFitness on unseen proteins (Figure S2b) by 10-fold cross validation to test the truely zero-shot prediction behavior, where 80% of the proteins were assigned for training/validation while the remaining 20% unseen proteins were used for testing. Even in such a challenging case, GeoFitness still exhibits a considerable level of the mean Spearman correlation coefficient, which falls between 0.3 and 0.4, consistent with our previous testing in the extremely low amount of training data (*e.g*., data proportion = 1% in Figure S3).

### 2.2 Potential of inferring multi-mutational effects by GeoFitness

Albeit designed only for predicting the effects caused by single mutations, GeoFitness is, in principle, capable of inferring the multiple mutational effects in a sequential manner. Here, we take a double mutation as an example to illustrate the procedure (Figure S4): 1) feed the sequence and structure features of the wild-type protein into GeoFitness and pick out the fitness score of the first single mutation (relative to the wide-type) from the full fitness landscape; 2) generate the 3D structure caused by the first mutation using Fold X; 3) feed the sequence and structure features of the protein carrying the first mutation into GeoFitness to obtain the fitness score of the second mutation (relative to the first mutation); and 4) add the two relative fitness scores as an inference for the effect of the corresponding double mutation. The process could be repeated sequentially for the inference of multiple mutational effects, although the results may depend on the order of mutations carried out virtually.

We collected double mutations from 7 proteins in the MaveDB database (Table S3). Specifically, we randomly chose 1000 pieces of double mutational data for each candidate protein. All data were retained if the amount of available data was less than 1000 for a protein. We then conducted zero-shot inference of double mutational effects, for the GeoFitness model that was originally trained by single mutational data. As shown in Table S3, the fitness score of double mutations predicted by GeoFitness correlate unexpectedly well with experimental values. The mean Spearman correlation coefficient is 0.48 when averaged over proteins and 0.52 when averaged over all double mutational variants.

### 2.3 Anti-symmetry in GeoDDG/GeoDTm

In GeoDDG/GeoDTm, input features of the wild-type and mutant proteins are fed into the same set of neural networks (composed of the pre-trained module as well as the downstream geometric encoder and MLP/output layers) with shared weights, and then the difference between the network readouts of wide-type and mutant proteins is scaled to generate the predicted ΔΔ*G*/Δ*T*_*m*_ values (see Figure 2). Theoretically, the weight sharing and substraction of readouts jointly guarantee the anti-symmetry of predicted ΔΔ*G*/Δ*T*_*m*_, considering that the final result automatically changes sign when the input features of wild-type and mutant proteins are swapped.

To strictly ensure the anti-symmetry of prediction, the 2D features of the wild-type and mutant proteins should be extracted from their corresponding 3D structures, respectively. However, in practice, only one of the two structures (wild-type/mutant) is available, and the other structure (mutant/wild-type) should be generated *in silico* using the software Fold X. Unfortunately, the anti-symmetry is broken in this process, since the wild-type-to-mutant and mutant-to-wild-type structure generations by Fold X are not reversible.

To clearly elucidate this effect, we tested the anti-symmetric properties of GeoDDG predictions upon two sets of proteins, S^sym76^ and S2000^77^, where experimental structures are available for both mutant and wide-type sequences for each protein. As shown in Table S6, anti-symmetry is strictly preserved when the experimental structures are employed for feature generation, but is lost to a slight extent when Fold X is engaged to produce the counterpart structure from the given one.

Nevertheless, based on our preliminary results, the difference in structural deviation caused by Fold X when wild-type or mutant structures are used as the starting point indeed exists but is usually small. However, if the user demands strict anti-symmetry for the prediction results, it is recommended to use the same set of wild-type and mutant structures for feature generation rather than to automatically follow the Fold X pipeline.

### 2.4 Evaluation of GeoDDG on additional protein stability data from DMS

In a recent work^78^, a large amount of experimental DMS data related with protein stability have been released. Although the labels are not purely identical to ΔΔ*G*, we chose around 20 thousand single mutational data from this large-scale dataset, including 160 proteins with experimental structures, to benchmark the performance of GeoDDG in such a zero-shot prediction task.

As shown in Figure S6, GeoDDG predictions correlate fairly well with the experimental DMS data. Specifically, when averaged over proteins, the mean Spearman correlation coefficient is 0.63, a value very close to the same metric evaluated over the S669 testing set (see Table 1). This observation supports the robustness of our model. It is worth noting that the data labels in the DMS dataset were inferred based on proteolysis assays and thus could not truly represent the experimentally determined ΔΔ*G*. Hence, metrics based on absolute values, like MAE and RMSE, should be less concerned in performance evaluation.

### 2.5 Performance of GeoDDG in the prediction of multi-mutational effects

We evaluated the performance of GeoDDG in the zero-shot prediction of ΔΔ*G* caused by multiple mutations. Specifically, we first collected multiple mutation data consisting of 133 proteins with 792 pieces of double mutation data and 469 pieces of triple or higher-order mutation data and composed a testing dataset named as M1261 (see **Methods**). Next, we evaluated the prediction power on ΔΔ*G* of multiple mutations for the GeoDDG model that was trained with single mutational data only as well as two mainstream models, DynaMut2^79^ and DDMut^50^, which were originally trained with both single and multiple mutational data. As shown in Table S7, GeoDDG shows comparable performance to DDMut in predicting the effects of double, triple and higher-order mutations in evaluation metrics reflecting the correlation (*i.e*. Spearman and Pearson correlation coefficients), whereas both methods remarkably outperform DynaMut2.

Considering that ΔΔ*G* data caused by multiple mutations are completely absent in the training set of GeoDDG, its good performance in the prediction of such effects further corroborates the good generalizability of our GeoDDG model.

### 2.6 Necessity and significance of the loss function and geometric encoder in GeoDDG/GeoDTm

The ablation studies from Figures 5, 6, and 7 suggest that the conjugate loss function and the geometric encoder both introduce relatively modest improvement to the model performance of GeoDDG and GeoDTm. Here, we continue to analyze these two factors by taking GeoDDG-3D as an examplar studying target.

Our previous ablation studies were conducted with all factors sequentially applied, including not only the geometric encoder and the conjugate loss, but also data expansion and the pre-training strategy whose contributions are understandable and have been thoroughly discussed in the main text. Here, in order to focus on the loss function and the geometric encoder, we first retrained a baseline GeoDDG model without employing the data expansion and the pre-training strategy. Specifically, the pre-trained module (from GeoFitness) was removed and the model was trained using the S2648 dataset only. Next, we performed two control experiments to analyze the roles of the conjugate loss and the geometric encoder in the model optimization, respectively.

On the one hand, we replaced the conjugate loss by the pure MSE loss as a control to explore the contribution of SCC loss. Although value of the loss function could be minimized in both cases (Figure S8a), the model trained with the pure MSE loss clearly exhibits lower Spearman correlation coefficients on the validation set (Figure S8b), since this loss alone is unable to capture the relative ordering information between labels .

On the other hand, we substituted the geometric encoder in the baseline model by MLP layers with a similar amount of learnable parameters as a control to study the role of geometric encoder. As expected, the model purely composed of MLP layers shows weakened Spearman correlation coefficients on the validation set (Figure S8c), because the MLP layers alone are inferior in handling protein structural features.

In summary, the above experiments further support the necessity and significance of both the conjugate loss function and the geometric encoder in the model designed in this work.

## 3 Supplementary Tables

**Table S1.** The details of DMS datasets for GeoFitness training.

| Dataset name | Type | Detailed Type | Number |
| --- | --- | --- | --- |
| BRCA1_HUMAN_BRCT<sup>[1]</sup> | Binding ability | Protein interaction | 1,262 |
| B3VI55_LIPSTSTABLE<sup>[1]</sup> | Enzyme function | Transferase activity | 7,890 |
| RL401_YEAST_Fraser2016<sup>[1]</sup> | Cell process | Protein modifier | 1,253 |
| BG_STRSQ_hmmerbit<sup>[1]</sup> | Enzyme function | Hydrolase activity | 2,999 |
| BF520_env_Bloom2018<sup>[1]</sup> | Binding ability | Protein interaction | 12,577 |
| B3VI55_LIPST_Whitehead2015<sup>[1]</sup> | Enzyme function | Transferase activity | 7,631 |
| GAL4_YEAST_Shendure2015<sup>[1]</sup> | Binding ability | DNA binding | 1,195 |
| TIM_SULSO_b0<sup>[1]</sup> | Enzyme function | Lyase activity | 1,519 |
| POLG_HCVJF_Sun2014<sup>[1]</sup> | Binding ability | drug binding | 1,630 |
| HIS7_YEAST_Kondrashov2017<sup>[1]</sup> | Enzyme function | Lyase activity | 168 |
| MTH3_HAEAEESTABILIZED_Tawfik2015<sup>[1]</sup> | Enzyme function | Transferase activity | 1,777 |
| AMIE_PSEAE_Whitehead<sup>[1]</sup> | Enzyme function | Hydrolase activity | 6,227 |
| P84126_THETH_b0<sup>[1]</sup> | Enzyme function | Lyase activity | 1,519 |
| IF1_ECOLI_Kishony<sup>[1]</sup> | Binding ability | DNA binding | 1,367 |
| PA_FLU_Sun2015<sup>[1]</sup> | Enzyme function | Transferase activity | 1,820 |
| KKA2_KLEPN_Mikkelsen2014<sup>[1]</sup> | Enzyme function | Transferase activity | 4,960 |
| TIM_THEMA_b0<sup>[1]</sup> | Enzyme function | Lyase activity | 1,519 |
| BG505_env_Bloom2018<sup>[1]</sup> | Binding ability | Protein interaction | 12,729 |
| HG_FLU_Bloom2016<sup>[1]</sup> | Binding ability | Protein interaction | 10,715 |
| BRCA1_HUMAN_RING<sup>[1]</sup> | Enzyme function | Transferase activity | 575 |
| RASH_HUMAN_Kuriyan<sup>[1]</sup> | Binding ability | Protein interaction | 3,134 |
| 00000001-a-1<sup>[2]</sup> | Enzyme function | Transferase activity | 3,021 |
| 00000001-b-1<sup>[2]</sup> | Cell process | Protein modifier | 1,919 |
| 00000001-c-2<sup>[2]</sup> | Cell process | Calmodulin function | 2,831 |
| 00000001-d-2<sup>[2]</sup> | Enzyme function | Transferase activity | 4,617 |
| 00000002-a-2<sup>[2]</sup> | Binding ability | Protein interaction | 513 |
| 00000004-a-3<sup>[2]</sup> | Enzyme function | Transferase activity | 940 |
| 00000005-a-3<sup>[2]</sup> | Enzyme function | Lyase activity | 10,450 |
| 00000010-a-1<sup>[2]</sup> | Binding ability | RNA binding | 1,188 |
| 00000011-a-1<sup>[2]</sup> | Cell process | Chaperon function | 171 |
| 00000013-b-1<sup>[2]</sup> | Expression | Protein abundance | 3,689 |
| 00000035-a-3<sup>[2]</sup> | Enzyme function | Oxidoreductase activity | 16,853 |
| 00000036-a-2<sup>[2]</sup> | Binding ability | Protein interaction | 5,814 |
| 00000041-a-1<sup>[2]</sup> | Enzyme function | Transferase activity | 3,714 |
| 00000043-a-2<sup>[2]</sup> | Cell process | Receptor activation | 562 |
| 00000044-b-2<sup>[2]</sup> | Expression | Protein abundance | 3,798 |
| 00000045-h-1<sup>[2]</sup> | Cell process | Cellular toxicity | 2,599 |
| 00000046-a-2<sup>[2]</sup> | Cell process | Resistance to ubiquitination | 467 |
| 00000047-a-1<sup>[2]</sup> | Expression | Cell surface protein abundance | 6,639 |

**Table S1. Continued from previous page**
| **Dataset name** | **Type** | **Detailed Type** | **Number** |
| --- | --- | --- | --- |
| 00000048-a-1<sup>[2]</sup> | Expression | Cell surface protein abundance | 6,645 |
| 00000049-a-7<sup>[2]</sup> | Enzyme function | Oxidoreductase activity | 11,357 |
| 00000050-a-1<sup>[2]</sup> | Cell process | DNA repair | 16,749 |
| 00000055-a-1<sup>[2]</sup> | Expression | Protein abundance | 2,922 |
| 00000059-a-1<sup>[2]</sup> | Binding ability | DNA binding | 2,967 |
| 00000060-a-1<sup>[2]</sup> | Cell process | Cellular toxicity | 1,342 |
| 00000061-a-1<sup>[2]</sup> | Binding ability | Protein interaction | 297 |
| 00000063-b-1<sup>[2]</sup> | Enzyme function | Oxidoreductase activity | 2,527 |
| 00000064-a-1<sup>[2]</sup> | Binding ability | Protein interaction | 1,870 |
| 00000065-a-1<sup>[2]</sup> | Cell process | GDP-GTP exchange | 2,997 |
| 00000066-a-1<sup>[2]</sup> | Cell process | Synaptic transmission | 4,365 |
| 00000067-a-1<sup>[2]</sup> | Enzyme function | Oxidoreductase activity | 3,756 |
| 00000068-a-1<sup>[2]</sup> | Cell process | Cell cycle (p53) | 7,467 |
| 00000069-a-1<sup>[2]</sup> | Binding ability | Protein interaction | 2,223 |
| 00000071-a-1<sup>[2]</sup> | Enzyme function | Hydrolase activity | 2,404 |
| 00000072-a-1<sup>[2]</sup> | Binding ability | Protein interaction | 666 |
| 00000073-c-1<sup>[2]</sup> | Binding ability | Substrate binding | 5,023 |
| 00000074-a-1<sup>[2]</sup> | Cell process | Chaperon function | 4,323 |
| 00000075-a-1<sup>[2]</sup> | Binding ability | Protein interaction | 9,576 |
| 00000076-a-1<sup>[2]</sup> | Cell process | Subcellular localization | 507 |
| 00000077-a-1<sup>[2]</sup> | Binding ability | Protein interaction | 2,986 |
| 00000077-b-1<sup>[2]</sup> | Binding ability | Protein interaction | 2,777 |
| 00000078-b-1<sup>[2]</sup> | Expression | Protein abundance | 2,695 |
| 00000079-a-1<sup>[2]</sup> | Expression | Protein abundance | 4,840 |
| 00000080-a-2<sup>[2]</sup> | Fluorescence | Fluorescence intensity | 1,084 |
| 00000084-a-1<sup>[2]</sup> | Cell process | Cellular toxicity | 1,176 |
| 00000087-a-1<sup>[2]</sup> | Enzyme function | Transferase activity | 9,462 |
| 00000090-b-1<sup>[2]</sup> | Binding ability | Protein interaction | 1,576 |
| 00000094-a-3<sup>[2]</sup> | Cell process | Channel function | 3,319 |
| 00000095-a-1<sup>[2]</sup> | Enzyme function | Oxidoreductase activity | 6,142 |
| 00000096-a-1<sup>[2]</sup> | Enzyme function | Transferase activity | 8,570 |
| 00000099-a-1<sup>[2]</sup> | Expression | Cell surface protein abundance | 165 |
| 00000102-0-1<sup>[2]</sup> | Expression | Protein abundance | 5,083 |
| 00000103-d-1<sup>[2]</sup> | Enzyme function | Transferase activity | 6,810 |

The superscript in the dataset name represents the source of the data, where [1] denotes data from DeepSequence and [2] denotes data from MaveDB.

**Table S2.** Features adopted by the GeoStab-suite.

| Feature | Shape | Source or meaning |
| --- | --- | --- |
| 1D information | [$L$, 1280] | Calculated by ESM-2 (esm2_t33_650M_UR50D) |
| Physicochemical properties | [$L$, 7] | Including steric parameters, hydrophobicity, volume, polarizability, isoelectric point, helix probability and sheet probability |
| pH&temp | 2 | pH and temperature |
| 2D information | [$L$, $L$, 7] | Calculated from atom coordinates |
| Esm1v-logits | [$L$, 20, 5] | Predicted by five ESM-1v models and used in the output layer of GeoFitness |
| pLDDT | [$L$, 1] | Reported by AlphaFold2 and applied to mask nodes and edges with low prediction quality in GeoDDG-Seq/GeoDTm-Seq |

$L$ refers to the number of residues of the target protein.

**Table S3.** Zero-shot inference of the double mutational effects by GeoFitness.

| Protein Name | Dataset Name | Single Mutation (Total Number) | Double Mutation (Total Number) | Single Mutation (Train Number) | Double Mutation (Test Number) | Double Mutation $\rho$ |
| --- | --- | --- | --- | --- | --- | --- |
| YAP1 | 00000002-a-2 | 513 | 19371 | 359 | 1000 | 0.63 |
| E4B | 00000004-a-3 | 940 | 52131 | 658 | 1000 | 0.52 |
| PAB1 | 00000010-a-1 | 1188 | 36522 | 832 | 1000 | 0.76 |
| TpoR | 00000043-a-2 | 562 | 199 | 393 | 199 | 0.22 |
| CD86 | 00000046-a-2 | 467 | 3710 | 327 | 1000 | 0.50 |
| p53 | 00000059-a-1 | 2967 | 1151 | 2077 | 1000 | 0.14 |
| GFP | 00000080-a-2 | 1084 | 12777 | 759 | 1000 | 0.60 |

The mean Spearman correlation coefficient ( $\rho$ ) is **0.48** when averaged over proteins and **0.52** when averaged over all double-mutation variants. GeoFitness was only trained by single mutational data but was tested on double mutational data. The dataset splitting strategy was the randomized split.

**Table S4.** Comparison of GeoDDG with exsiting models on the S461 dataset.

| Method | Total |  | Direct |  | Inverse |  | Antisymmetry |  |
| --- | --- | --- | --- | --- | --- | --- | --- | --- |
| | $\rho/r$ | RMSE/MAE | $\rho/r$ | RMSE/MAE | $\rho/r$ | RMSE/MAE | $r_{d-i}$ | $\langle\delta\rangle$ |
| *Sequence-based* |  |  |  |  |  |  |  |  |
| INPS-Seq | 0.79/0.77 | 1.11/0.82 | 0.55/0.56 | 1.10/0.82 | 0.54/0.55 | 1.12/0.83 | **-1.00** | 0.03 |
| ACDC-NN-Seq | 0.76/0.76 | 1.10/0.80 | 0.54/0.57 | 1.10/0.80 | 0.54/0.57 | 1.10/0.80 | **-1.00** | **0.00** |
| DDGun | 0.77/0.76 | 1.29/0.96 | 0.58/0.59 | 1.26/0.95 | 0.55/0.55 | 1.32/0.97 | -0.94 | 0.07 |
| I-Mutant3.0-Seq | 0.45/0.47 | 1.69/1.32 | 0.45/0.44 | 1.16/0.89 | 0.25/0.24 | 2.09/1.75 | -0.43 | 0.79 |
| MuPro | 0.43/0.47 | 1.79/1.42 | 0.37/0.40 | 1.17/0.91 | 0.27/0.29 | 2.24/1.93 | -0.31 | 0.97 |
| SAAFEC-Seq | 0.25/0.35 | 1.80/1.37 | 0.48/0.49 | 1.12/0.83 | 0.01/0.02 | 2.29/1.92 | -0.05 | 0.85 |
| GeoDDG-Seq | **0.83/0.82** | **0.98/0.73** | **0.68/0.69** | **0.98/0.73** | **0.68/0.69** | **0.98/0.73** | **-1.00** | **0.00** |
| *Structure-based* |  |  |  |  |  |  |  |  |
| DDMut | 0.79/0.79 | 1.05/0.80 | 0.60/0.62 | 1.03/0.77 | 0.57/0.58 | 1.08/0.82 | -0.94 | 0.13 |
| PremPS | 0.81/**0.81** | **1.01/0.76** | 0.60/0.63 | 1.03/0.79 | 0.58/0.63 | **0.98/0.73** | -0.86 | 0.20 |
| ACDC-NN | 0.78/0.78 | 1.07/0.77 | 0.57/0.61 | 1.06/0.77 | 0.56/0.60 | 1.07/0.78 | -0.98 | 0.04 |
| INPS3D | 0.71/0.69 | 1.27/0.93 | 0.60/0.62 | **1.01/0.75** | 0.39/0.35 | 1.48/1.12 | -0.49 | 0.48 |
| DDGun3D | 0.72/0.76 | 1.14/0.83 | 0.58/0.64 | 1.10/0.80 | 0.55/0.59 | 1.17/0.85 | -0.96 | 0.10 |
| DynaMut | 0.59/0.63 | 1.32/0.98 | 0.44/0.50 | 1.27/0.96 | 0.43/0.44 | 1.36/1.01 | -0.61 | 0.29 |
| ThermoNet | 0.60/0.67 | 1.27/0.97 | 0.49/0.56 | 1.23/0.93 | 0.43/0.52 | 1.31/1.01 | -0.86 | 0.16 |
| PopMusic | 0.56/0.60 | 1.51/1.14 | 0.61/0.62 | **1.01/0.76** | 0.25/0.29 | 1.89/1.52 | -0.30 | 0.69 |
| FoldX | 0.53/0.44 | 2.00/1.26 | 0.35/0.22 | 2.23/1.38 | 0.45/0.41 | 1.74/1.14 | -0.22 | 0.71 |
| MAESTRO | 0.48/0.54 | 1.54/1.16 | 0.60/0.63 | 1.04/0.78 | 0.17/0.20 | 1.92/1.53 | -0.22 | 0.68 |
| DUET | 0.46/0.52 | 1.58/1.18 | 0.57/0.59 | 1.06/0.77 | 0.15/0.18 | 1.97/1.59 | -0.15 | 0.72 |
| mCSM | 0.40/0.46 | 1.71/1.31 | 0.51/0.53 | 1.07/0.80 | 0.12/0.18 | 2.17/1.81 | -0.09 | 0.85 |
| I-Mutant3.0 | 0.34/0.41 | 1.74/1.33 | 0.48/0.49 | 1.12/0.83 | 0.11/0.13 | 2.19/1.83 | -0.04 | 0.81 |
| SDM | 0.33/0.41 | 1.69/1.26 | 0.51/0.56 | 1.33/1.01 | 0.11/0.11 | 1.98/1.51 | -0.35 | 0.52 |
| GeoDDG-3D | **0.83/0.81** | **1.01/0.72** | **0.67/0.67** | **1.01/0.72** | **0.67/0.67** | **1.01/0.72** | **-1.00** | **0.00** |

The winner in each metric is highlighted in bold. "Direct" refers to the mutations from the wild type to mutant, "Inverse" means the corresponding mutations in the reverse direction, and "Total" stands for the combination of the above categories. Here, $\rho$ and $r$ denote Spearman and Pearson correlation coefficients, respectively. Details of all metrics are described in Methods. The raw results of the other models except DDMut<sup>50</sup> are taken from a prior work<sup>52</sup>.

**Table S5.** Comparison of GeoDDG-LE with exsiting models on the S783 dataset.

| Method | Total |  | Direct |  | Inverse |  | Antisymmetry |  |
| --- | --- | --- | --- | --- | --- | --- | --- | --- |
| | $\rho/r$ | RMSE/MAE | $\rho/r$ | RMSE/MAE | $\rho/r$ | RMSE/MAE | $r_{d-i}$ | $\langle\delta\rangle$ |
| *Sequence-based* |  |  |  |  |  |  |  |  |
| INPS-Seq |  |  | 0.56/0.59 | 1.68/1.15 |  |  |  |  |
| GeoDDG-LE-Seq | **0.80/0.82** | **1.51/1.01** | **0.72/0.73** | **1.51/1.01** | **0.72/0.73** | **1.51/1.01** | **-1.00** | **0.00** |
| *Structure-based* |  |  |  |  |  |  |  |  |
| PremPS |  |  | 0.66/**0.72** | **1.51/1.05** |  |  |  |  |
| PoPMuSiC |  |  | 0.60/0.67 | **1.51/1.05** |  |  |  |  |
| INPS3D |  |  | 0.59/0.65 | 1.58/1.07 |  |  |  |  |
| FoldX |  |  | 0.60/0.57 | 1.99/1.26 |  |  |  |  |
| mCSM |  |  | 0.50/0.58 | 1.65/1.17 |  |  |  |  |
| GeoDDG-LE-3D | **0.78/0.80** | **1.55/1.05** | **0.71/0.72** | **1.55/1.05** | **0.71/0.72** | **1.55/1.05** | **-1.00** | **0.00** |

The winner in each metric is highlighted in bold. "Direct" refers to the mutations from the wild type to mutant, "Inverse" means the corresponding mutations in the reverse direction, and "Total" stands for the combination of the above categories. Here, $\rho$ and $r$ denote Spearman and Pearson correlation coefficients, respectively. Details of all metrics are described in **Methods**. The raw results of the other models are taken from a prior work<sup>49</sup>. Blank region means that the corresponding result is unavailable.

**Table S6.** Anti-symmetric properties of GeoDDG tested on the S_sym_ and S2000 sets.

| Table S6. Anti-symmetric properties of GeoDDG tested on the $S^{\text{sym}}$ and S2000 sets | | | | |
| --- | --- | --- | --- | --- |
| Dataset | Mutant structure built from Fold X |  | Both from experimental structures |  |
| | $r_{d-i}$ | $\langle \delta \rangle$ | $r_{d-i}$ | $\langle \delta \rangle$ |
| $S^{\text{sym}}$ | -0.999 | 0.013 | -1.000 | 0.000 |
| S2000 | -0.999 | 0.012 | -1.000 | 0.000 |

**Table S7.**
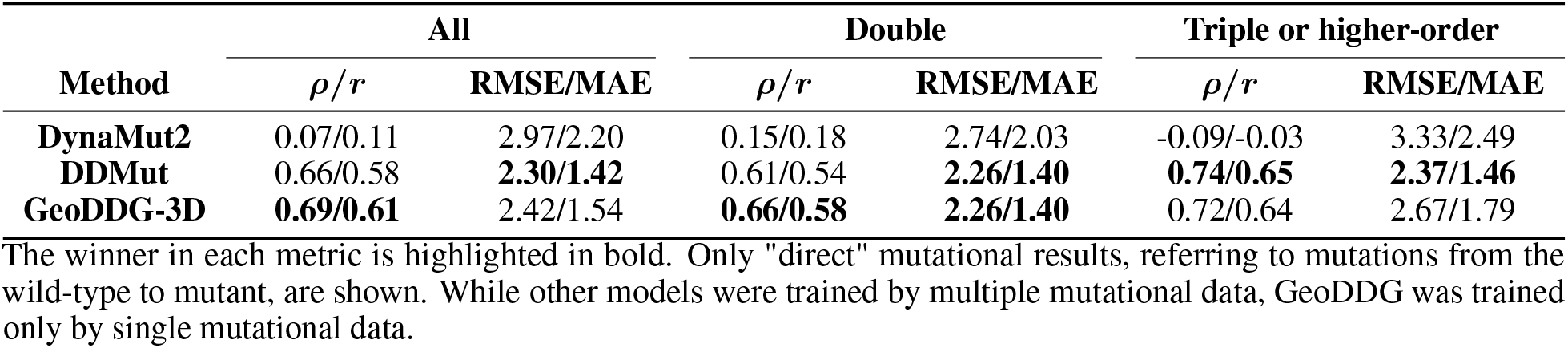
Comparison of GeoDDG with exsiting models on the prediction of multiple mutational effects.

| Method | All |  | Double |  | Triple or higher-order |  |
| --- | --- | --- | --- | --- | --- | --- |
| | $\rho/r$ | RMSE/MAE | $\rho/r$ | RMSE/MAE | $\rho/r$ | RMSE/MAE |
| **DynaMut2** | 0.07/0.11 | 2.97/2.20 | 0.15/0.18 | 2.74/2.03 | -0.09/-0.03 | 3.33/2.49 |
| **DDMut** | 0.66/0.58 | **2.30/1.42** | 0.61/0.54 | **2.26/1.40** | **0.74/0.65** | **2.37/1.46** |
| **GeoDDG-3D** | **0.69/0.61** | 2.42/1.54 | **0.66/0.58** | **2.26/1.40** | 0.72/0.64 | 2.67/1.79 |

The winner in each metric is highlighted in bold. Only "direct" mutational results, referring to mutations from the wild-type to mutant, are shown. While other models were trained by multiple mutational data, GeoDDG was trained only by single mutational data.

**Table S8.** Hyper-parameter search and selection for GeoDDG.

| *node_dim* | *edge_dim* | *num_head* | *num_layer* | number of<br>parameters<br>($\times 10^6$) | Spearman $\rho$ in<br>validation |
| --- | --- | --- | --- | --- | --- |
| 32 | 16 | 4 | 1 | 2.18 | 0.723 |
| 32 | 32 | 4 | 1 | 2.19 | 0.738 |
| 32 | 16 | 8 | 1 | 2.18 | 0.732 |
| 32 | 32 | 8 | 1 | 2.20 | 0.742 |
| 32 | 16 | 4 | 2 | 2.19 | 0.727 |
| 32 | 32 | 4 | 2 | 2.21 | 0.737 |
| 32 | 16 | 8 | 2 | 2.20 | 0.745 |
| 32 | 32 | 8 | 2 | 2.24 | 0.739 |
| 32 | 16 | 4 | 3 | 2.21 | 0.743 |
| 32 | 32 | 4 | 3 | 2.24 | 0.741 |
| 32 | 16 | 8 | 3 | 2.22 | 0.732 |
| 32 | 32 | 8 | 3 | 2.27 | 0.750 |
| 64 | 16 | 4 | 1 | 2.23 | 0.747 |
| 64 | 32 | 4 | 1 | 2.25 | 0.747 |
| 64 | 16 | 8 | 1 | 2.24 | 0.748 |
| 64 | 32 | 8 | 1 | 2.27 | 0.750 |
| 64 | 16 | 4 | 2 | 2.27 | 0.750 |
| 64 | 32 | 4 | 2 | 2.31 | 0.742 |
| 64 | 16 | 8 | 2 | 2.29 | 0.733 |
| 64 | 32 | 8 | 2 | 2.34 | 0.741 |
| 64 | 16 | 4 | 3 | 2.32 | 0.742 |
| 64 | 32 | 4 | 3 | 2.36 | 0.742 |
| 64 | 16 | 8 | 3 | 2.34 | 0.741 |
| 64 | 32 | 8 | 3 | 2.42 | 0.755 |
| 128 | 16 | 4 | 1 | 2.43 | 0.757 |
| 128 | 32 | 4 | 1 | 2.46 | 0.756 |
| 128 | 16 | 8 | 1 | 2.45 | 0.748 |
| 128 | 32 | 8 | 1 | 2.49 | 0.752 |
| 128 | 16 | 4 | 2 | 2.57 | 0.759 |
| 128 | 32 | 4 | 2 | 2.62 | 0.751 |
| 128 | 16 | 8 | 2 | 2.60 | 0.755 |
| 128 | 32 | 8 | 2 | 2.69 | 0.754 |
| 128 | 16 | 4 | 3 | 2.70 | 0.754 |
| 128 | 32 | 4 | 3 | 2.78 | 0.757 |
| 128 | 16 | 8 | 3 | 2.76 | 0.753 |
| 128 | 32 | 8 | 3 | 2.88 | 0.748 |
| **256** | **16** | **4** | **1** | **3.16** | **0.764** |
| 256 | 32 | 4 | 1 | 3.20 | 0.755 |
| 256 | 16 | 8 | 1 | 3.19 | 0.753 |
| 256 | 32 | 8 | 1 | 3.27 | 0.710 |
| 256 | 16 | 4 | 2 | 3.66 | 0.751 |
| 256 | 32 | 4 | 2 | 3.74 | 0.753 |
| 256 | 16 | 8 | 2 | 3.73 | 0.752 |
| 256 | 32 | 8 | 2 | 3.88 | 0.747 |
| 256 | 16 | 4 | 3 | 4.16 | 0.754 |
| 256 | 32 | 4 | 3 | 4.28 | 0.744 |
| 256 | 16 | 8 | 3 | 4.26 | 0.751 |
| 256 | 32 | 8 | 3 | 4.48 | 0.749 |
| 256 | 32 | 8 | 3 | 4.27 | 0.749 |

Here, *node\_dim* and *edge\_dim* denote the dimensions of the node and edge embeddings, respectively, *num\_head* refers to the number of attention heads, whereas *num\_layer* stands for the number of GAT layers in the geometric encoder (downstream to the pre-trained module). The set of hyper-parameters with the best performance is highlighted in bold.

**Table S9.** Impact of initialization methods and random seeds on the model performance of GeoDDG.

| | Details | Spearman $\rho$ in validation |
| --- | --- | --- |
| **Initialization Method** | default_initialization | **0.743** |
|  | kaiming_norm | 0.693 |
|  | xavier_normal | 0.711 |
|  | xavier_uniform | 0.690 |
|  | orthogonal | 0.708 |
| **Random Seed** | 50 repetitions | $0.743 \pm 0.004$ |

## 4 Supplementary Figures

**Figure S1.**
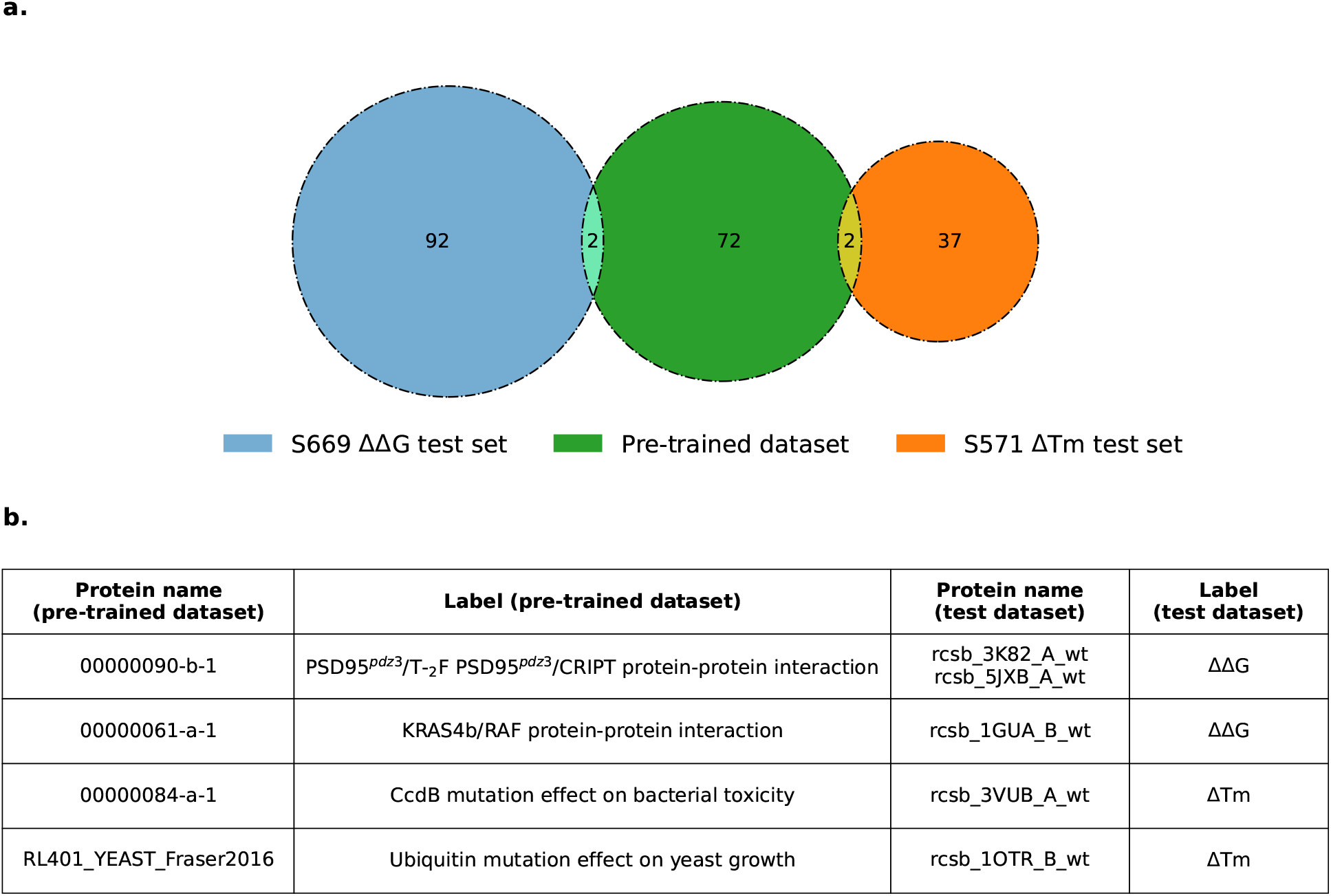
Lack of overlap between the datasets used for model pre-training and benchmark testing. **a)** The DMS dataset adopted for pre-training the GeoFitness model contains data collected from 74 proteins in total, among which 2 protein entries exhibit sequence identity > 40% with the proteins adopted by S669 and S571, the test datasets for ΔΔ*G* and Δ*T*_*m*_ predictions, respectively. **b)** The labels of the shared proteins in the pre-training dataset are, however, totally different from the metrics of protein stability, say ΔΔ*G* and Δ*T*_*m*_. Hence, there is no overlap between the datasets for model pre-training and benchmark testing, thereby precluding the possibility of information leaking during performance evaluation.

**Figure S2.**
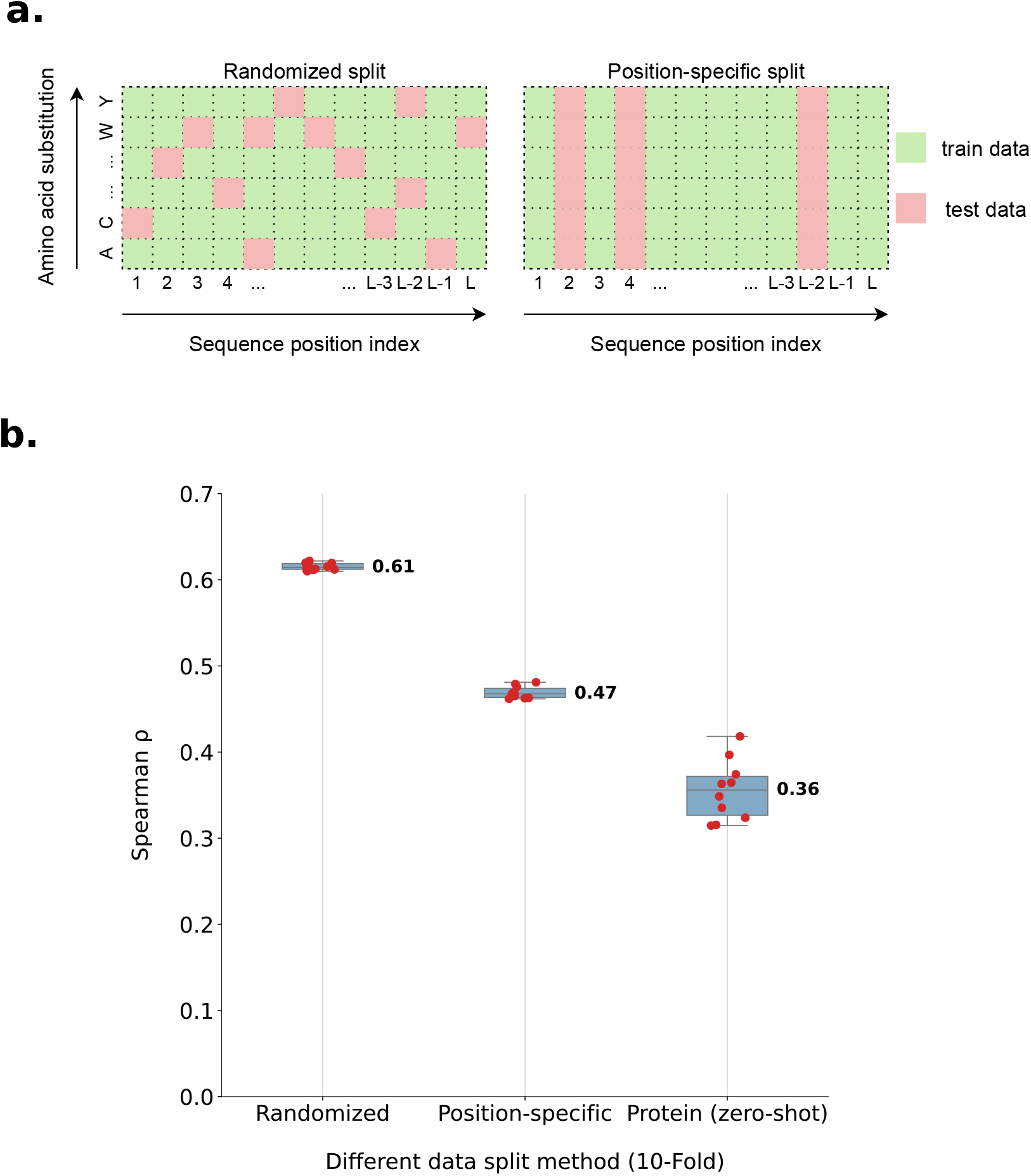
Three strategies of dataset splitting for the training of GeoFitness and their corresponding effects. **a)** Graphical interpretation of randomized split and position-specific split. The data “masked” for testing (colored pink) are chosen completely randomly in the randomized split strategy, but is applied column-wise in the position-specific split strategy. **b)** GeoFitness is evaluated with 10-fold cross validation using three different dataset splitting strategies. “Protein (zero-shot)” denotes the third splitting strategy at the protein level, with 80% proteins used for training/validation and 20% unseen proteins used for testing. Each red dot refers to the result of one individual experiment in the 10-fold cross validation, *i.e*. the mean Spearman *ρ* of all proteins. The center line of each box plot shows the median of the validation results with the value marked aside. The box limits correspond to the upper and lower quartiles, whereas the whiskers extend to points that lie within 1.5 inter-quartile range of the lower and upper quartiles.

**Figure S3.**
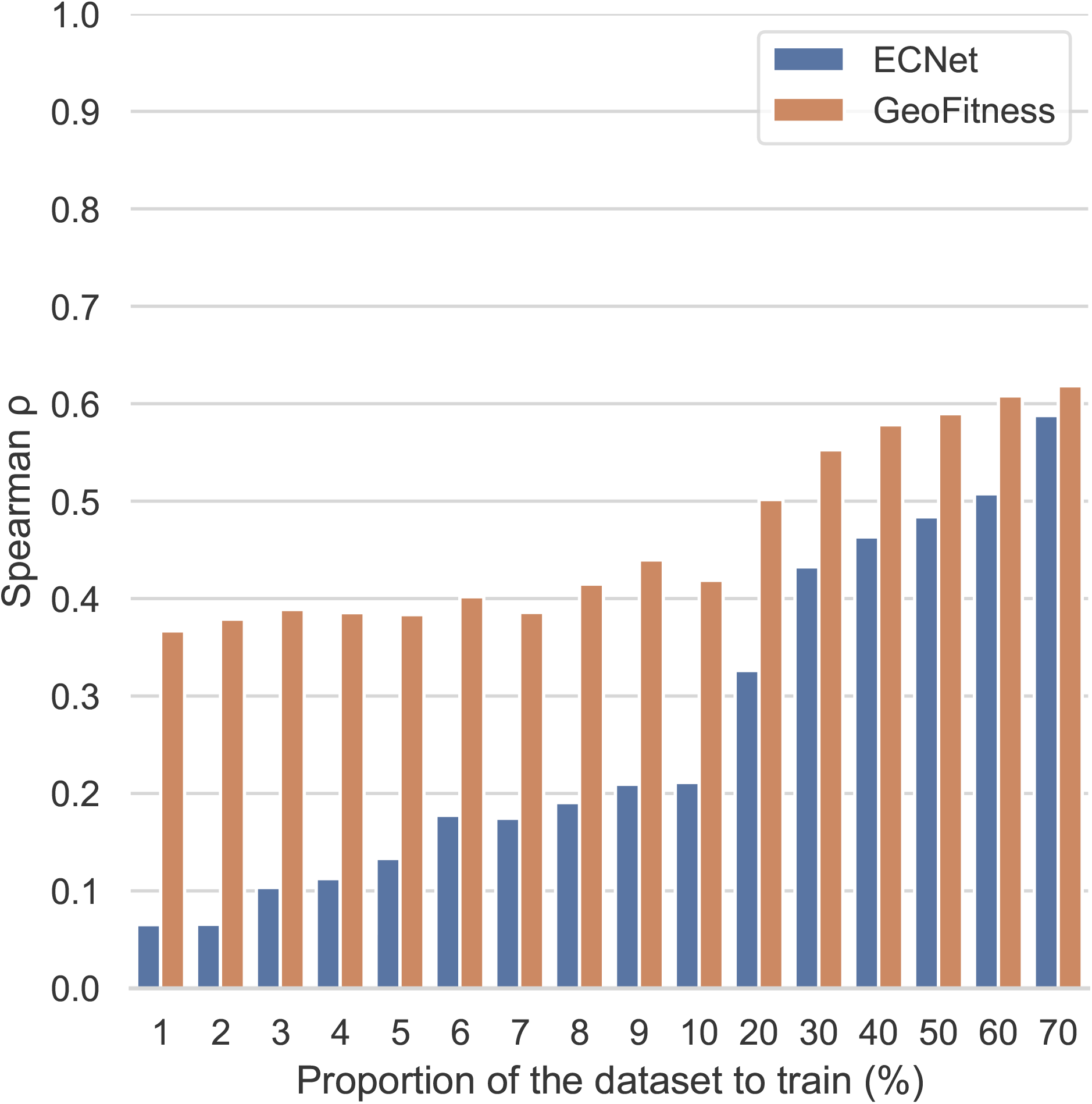
Comparison of GeoFitness and ECNet when using various proportion of data in the dataset for model training. The models retrained using different proportion of data in the original training set are evaluated on the original test set. Notably, the original training set contains 70% of the data in the whole DMS dataset, which corresponds to the maximum value on the horizontal axis in the bar chart.

**Figure S4.**
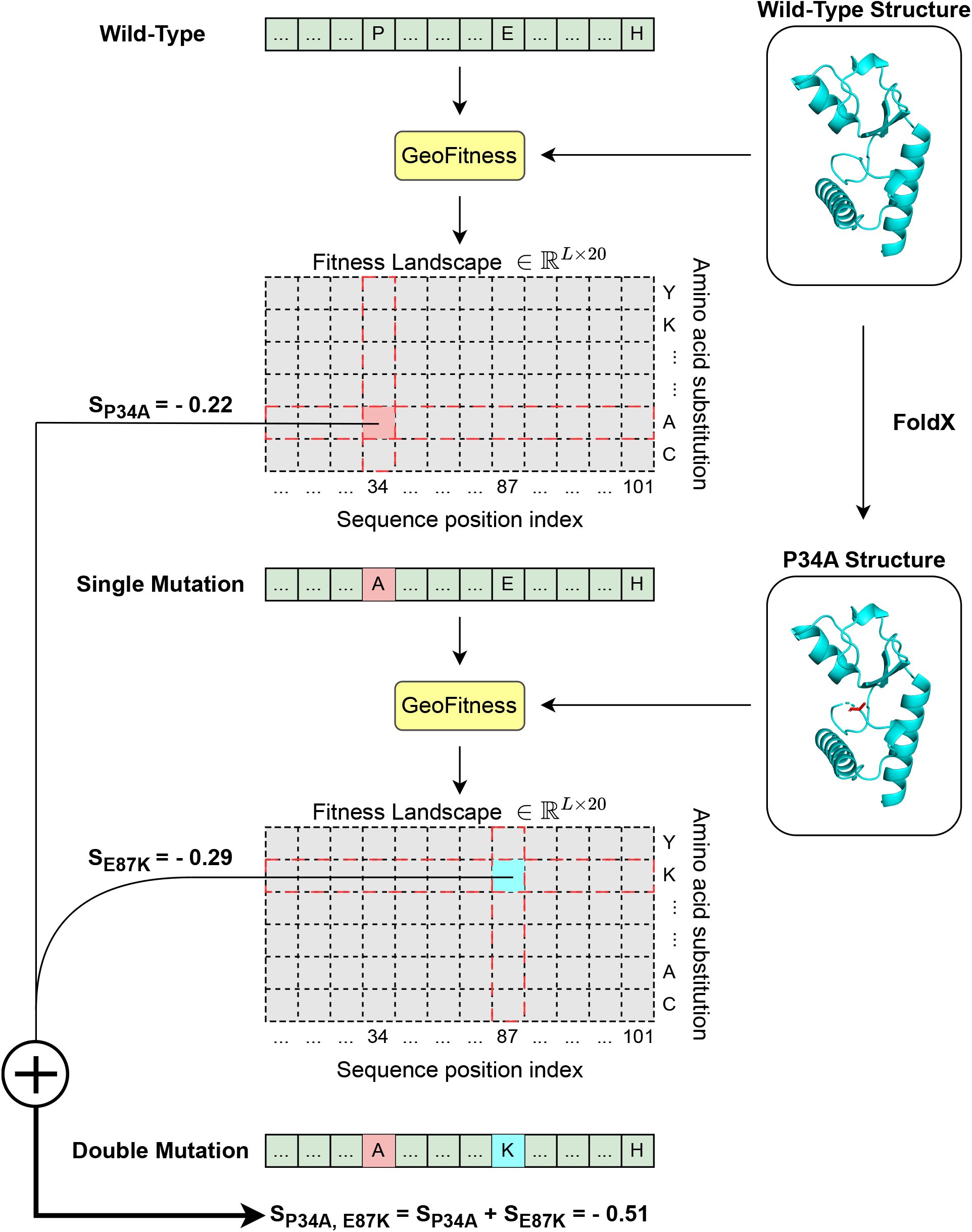
Inferring the multiple mutational effects by GeoFitness. Though designed only for single mutation prediction, GeoFitness has the potential to predict multiple mutational effects using the pipeline. Here, we show an example for the inference of double mutational effects. “S” in the figure means the fitness score predicted by GeoFitness. The process could be repeated for inferring the effects of more mutations.

**Figure S5.**
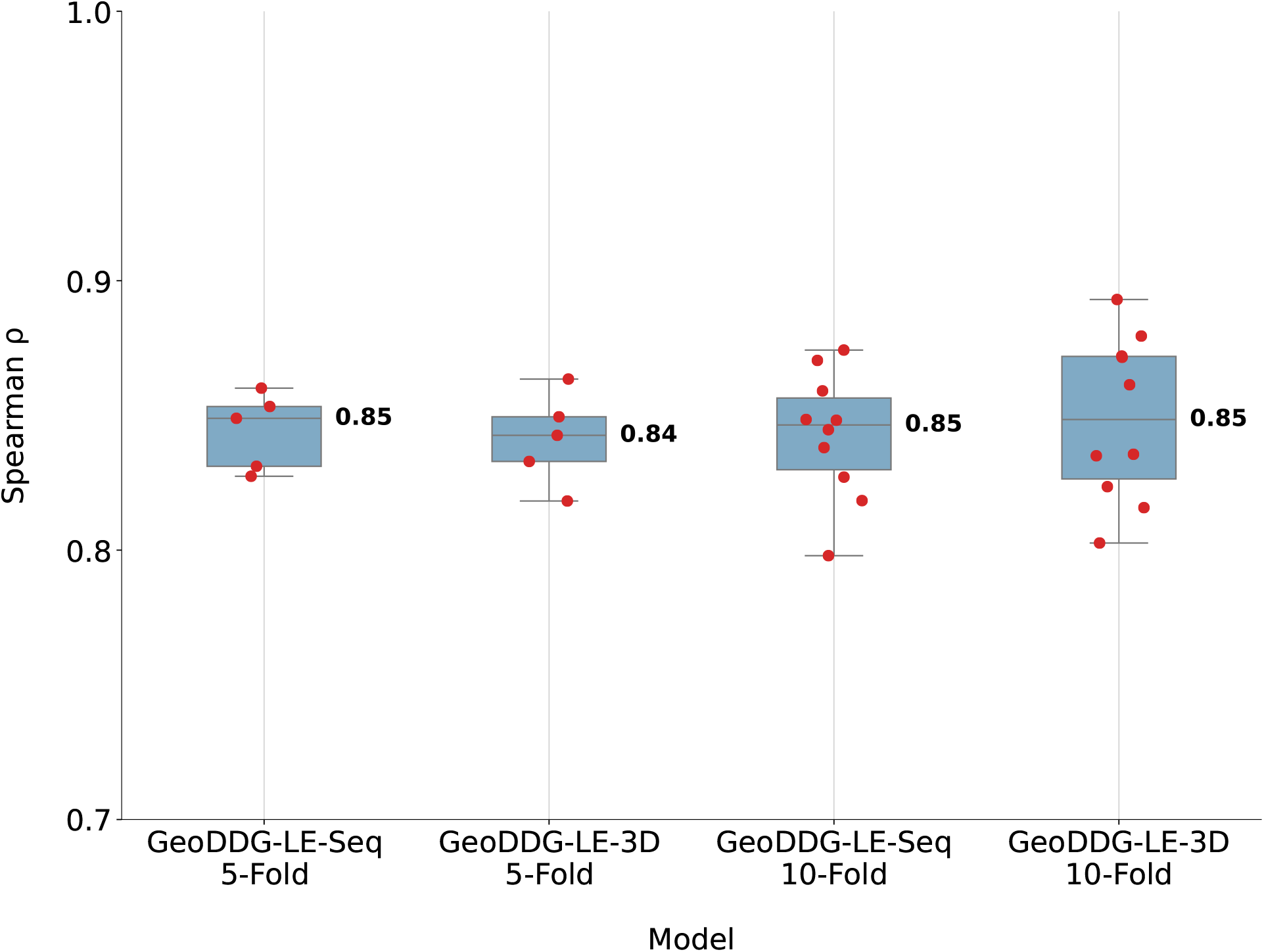
5-fold and 10-fold cross validation for GeoDDG-LE on the S2648 dataset. The “Seq” and “3D” versions of GeoDDG-LE are evaluated on S2648 with both 5-fold and 10-fold cross validations. Each red dot refers to an individual result of the cross validation. The center line of each box plot shows the median of the validation results with the value marked aside. The box limits correspond to the upper and lower quartiles, whereas the whiskers extend to points that lie within 1.5 inter-quartile range of the lower and upper quartiles.

**Figure S6.**
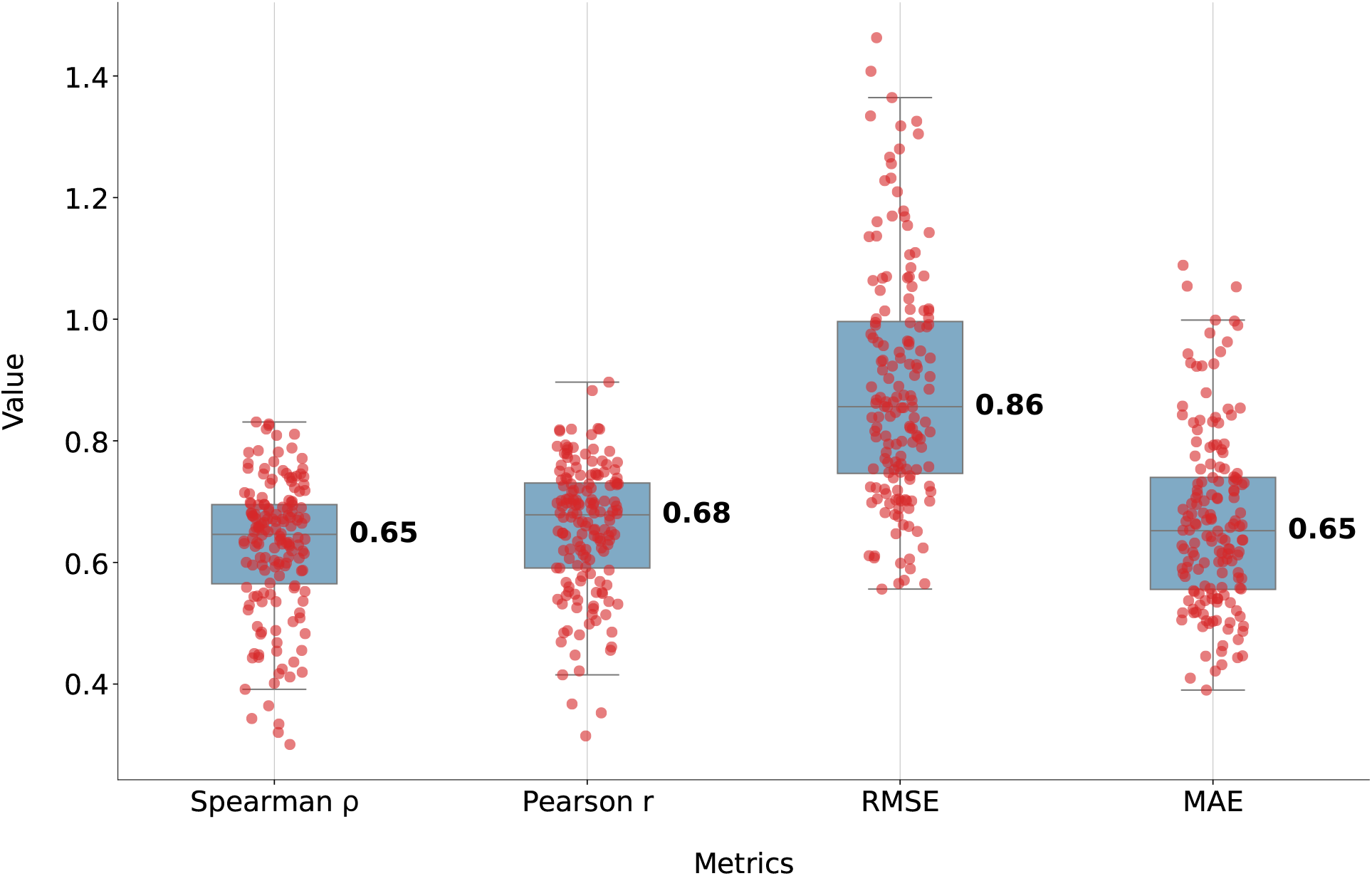
Evaluation of GeoDDG on a large scale DMS dataset whose labels are related with protein stability. GeoDDG was benchmarked on a large scale DMS dataset involving 160 proteins. Since each protein has at least 51 pieces of mutation data, the Spearman correlation coefficient *ρ*, Pearson correlation coefficient *r*, RMSE and MAE metrics can be calculated and averaged over each protein. Each red dot refers to the corresponding metric for an individual protein. The center line of each box plot shows the median of the metric with the value marked aside. The box limits correspond to the upper and lower quartiles, whereas the whiskers extend to points that lie within 1.5 inter-quartile range of the lower and upper quartiles.

**Figure S7.**
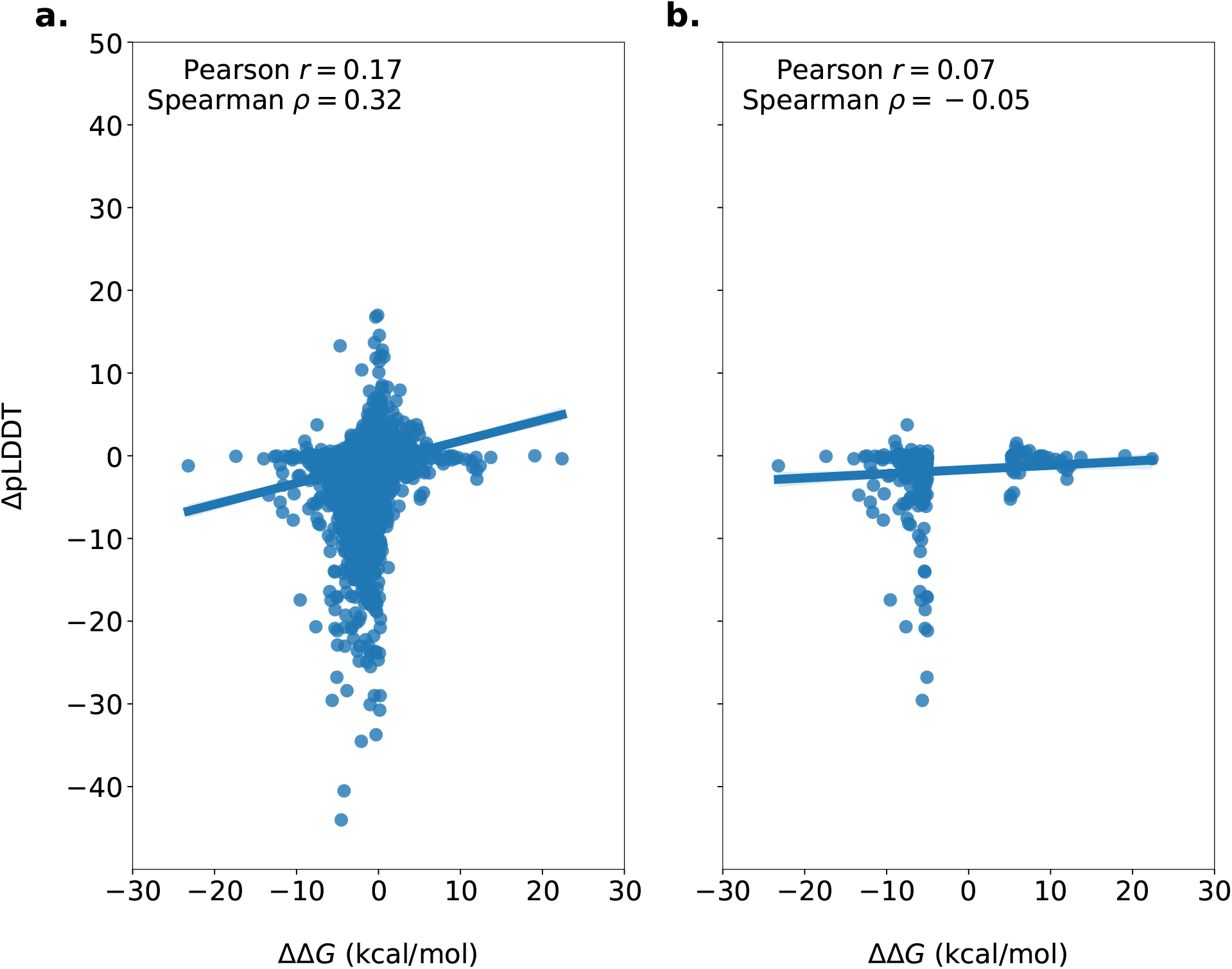
The correlation between ΔΔ*G* from experiments and ΔpLDDT by AlphaFold2. For each individual protein variant (shown as a dot in the figures), we calculated the difference of its Δ*G* and pLDDT (reported by AlphaFold2) with the corresponding value of the wide-type protein to evaluate the mutational effects on protein stability and AlphaFold2 prediction. **a)** All protein variants are considered. **b)** Only protein variants of high significance (*i.e*. |ΔΔ*G*| > 5 kcal/mol) are considered.

**Figure S8.**
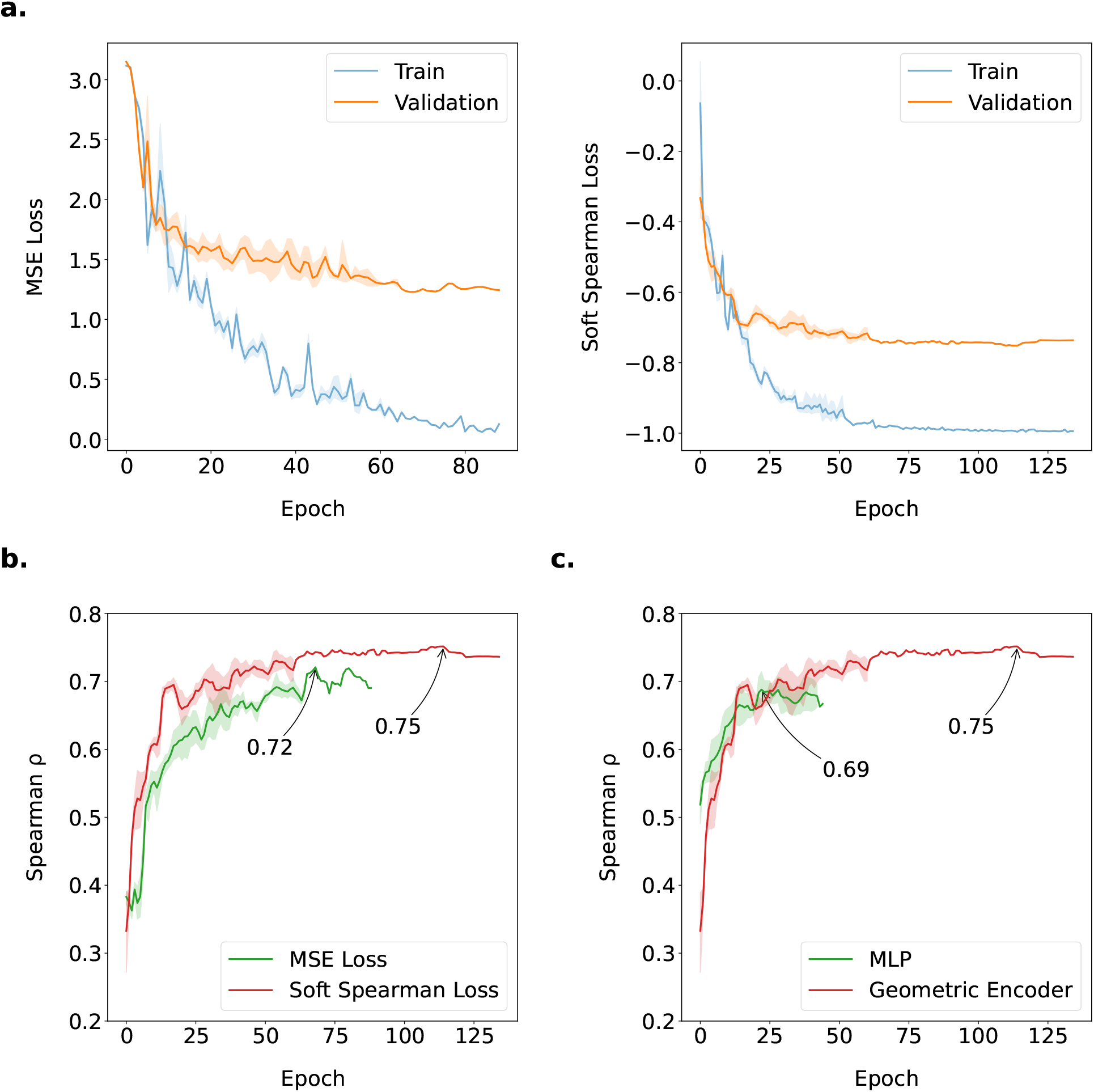
Exploration of the necessity of the conjugate loss function and the geometric encoder. **a)** The evolution of the training and validation losses in the training process when different loss functions are applied for model optimization: the MSE loss (left panel) vs. the conjugate loss (right panel). **b)** Evolution of the Spearman correlation coefficient *ρ* between inference and label on the validation set in the training process when different loss functions are used: the MSE loss (green) vs. the conjugate loss (red). **c)** Evolution of the Spearman correlation coefficient *ρ* between inference and label on the validation set in the training process when different model architectures are adopted: MLP (green) vs. geometric encoder (red). The error band is calculated across the 50 repetitions with different random seeds.

## References

1. Riesselman, A. J., Ingraham, J. B. & Marks, D. S. Deep generative models of genetic variation capture the effects of mutations. Nature Methods 15, 816–822. 10.1038/s41592-018-0138-4 (2018).

2. Coluzza, I. Computational protein design: a review. Journal of Physics: Condensed Matter 29, 143001. 10.1088/1361-648x/aa5c76 (2017).

3. Dahiyat, B. I. & Mayo, S. L. De Novo Protein Design: Fully Automated Sequence Selection. Science 278, 82–87. 10.1126/science.278.5335.82 (1997).

4. Tokuriki, N. & Tawfik, D. S. Stability effects of mutations and protein evolvability. Current Opinion in Structural Biology 19, 596–604. 10.1016/j.sbi.2009.08.003 (2009).

5. Pucci, F., Bourgeas, R. & Rooman, M. High-quality Thermodynamic Data on the Stability Changes of Proteins Upon Single-site Mutations. Journal of Physical and Chemical Reference Data 45, 023104. 10.1063/1.4947493 (2016).

6. Louis, B. B. V. & Abriata, L. A. Reviewing Challenges of Predicting Protein Melting Temperature Change Upon Mutation Through the Full Analysis of a Highly Detailed Dataset with High-Resolution Structures. Molecular Biotechnology 63, 863–884. 10.1007/s12033-021-00349-0 (2021).

7. Yeoman, C. J., Han, Y., Dodd, D., et al. in Advances in Applied Microbiology 1–55 (Elsevier, 2010). 10.1016/s0065-2164(10)70001-0.

8. Kopanos, C., Tsiolkas, V., Kouris, A., et al. VarSome: the human genomic variant search engine. Bioinformatics 35 (ed Wren, J.) 1978–1980. 10.1093/bioinformatics/bty897 (2018).

9. Fowler, D. M. & Fields, S. Deep mutational scanning: a new style of protein science. Nature Methods 11, 801–807. 10.1038/nmeth.3027 (2014).

10. McGuinness, K. N., Pan, W., Sheridan, R. P., et al. Role of simple descriptors and applicability domain in predicting change in protein thermostability. PLOS ONE 13 (ed Permyakov, E. A.) e0203819. 10.1371/journal.pone.0203819 (2018).

11. Wu, Z., Kan, S. B. J., Lewis, R. D., et al. Machine learning-assisted directed protein evolution with combinatorial libraries. Proceedings of the National Academy of Sciences 116, 8852–8858. 10.1073/pnas.1901979116 (2019).

12. Yang, K. K., Wu, Z. & Arnold, F. H. Machine-learning-guided directed evolution for protein engineering. Nature Methods 16, 687–694. 10.1038/s41592-019-0496-6 (2019).

13. Alley, E. C., Khimulya, G., Biswas, S., et al. Unified rational protein engineering with sequence-based deep representation learning. Nature Methods 16, 1315–1322. 10.1038/s41592-019-0598-1 (2019).

14. Amrein, B. A., Steffen-Munsberg, F., Szeler, I., et al. CADEE: Computer-Aided Directed Evolution of Enzymes. IUCrJ 4, 50–64. 10.1107/s2052252516018017 (2017).

15. Bedbrook, C. N., Yang, K. K., Rice, A. J., et al. Machine learning to design integral membrane channelrhodopsins for efficient eukaryotic expression and plasma membrane localization. PLOS Computational Biology 13 (ed Maranas, C. D.) e1005786. 10.1371/journal.pcbi.1005786 (2017).

16. Bedbrook, C. N., Yang, K. K., Robinson, J. E., et al. Machine learning-guided channelrhodopsin engineering enables minimally invasive optogenetics. Nature Methods 16, 1176–1184. 10.1038/s41592-019-0583-8 (2019).

17. Biswas, S., Khimulya, G., Alley, E. C., et al. Low-N protein engineering with data-efficient deep learning. Nature Methods 18, 389–396. 10.1038/s41592-021-01100-y (2021).

18. Hopf, T. A., Ingraham, J. B., Poelwijk, F. J., et al. Mutation effects predicted from sequence co-variation. Nature Biotechnology 35, 128–135. 10.1038/nbt.3769 (2017).

19. Khersonsky, O., Lipsh, R., Avizemer, Z., et al. Automated Design of Efficient and Functionally Diverse Enzyme Repertoires. Molecular Cell 72, 178–186.e5. 10.1016/j.molcel.2018.08.033 (2018).

20. Li, M., Kang, L., Xiong, Y., et al. SESNet: sequence-structure feature-integrated deep learning method for data-efficient protein engineering. Journal of Cheminformatics 15. 10.1186/s13321-023-00688-x (2023).

21. Luo, Y., Jiang, G., Yu, T., et al. ECNet is an evolutionary context-integrated deep learning framework for protein engineering. Nature Communications 12. 10.1038/s41467-021-25976-8 (2021).

22. Meier, J., Rao, R., Verkuil, R., et al. Language models enable zero-shot prediction of the effects of mutations on protein function in Advances in Neural Information Processing Systems (eds Ranzato, M., Beygelzimer, A., Dauphin, Y., et al.) 34 (Curran Associates, Inc., 2021), 29287–29303. https://proceedings.neurips.cc/paper_files/paper/2021/file/f51338d736f95dd42427296047067694-Paper.pdf.

23. Rao, R., Liu, J., Verkuil, R., et al. MSA Transformer. bioRxiv. 10.1101/2021.02.12.430858 (2021).

24. Rao, R., Bhattacharya, N., Thomas, N., et al. Evaluating Protein Transfer Learning with TAPE in Advances in Neural Information Processing Systems (eds Wallach, H., Larochelle, H., Beygelzimer, A., et al.) 32 (Curran Associates, Inc., 2019). https://proceedings.neurips.cc/paper_files/paper/2019/file/37f65c068b7723cd7809ee2d31d7861c-Paper.pdf.

25. Romero, P. A., Krause, A. & Arnold, F. H. Navigating the protein fitness landscape with Gaussian processes. Proceedings of the National Academy of Sciences 110. 10.1073/pnas.1215251110 (2012).

26. Russ, W. P., Figliuzzi, M., Stocker, C., et al. An evolution-based model for designing chorismate mutase enzymes. Science 369, 440–445. 10.1126/science.aba3304 (2020).

27. Wang, J., Lisanza, S., Juergens, D., et al. Scaffolding protein functional sites using deep learning. Science 377, 387–394. 10.1126/science.abn2100 (2022).

28. Mansoor, S., Baek, M., Juergens, D., et al. Accurate Mutation Effect Prediction using RoseTTAFold. bioRxiv. 10.1101/2022.11.04.515218 (2022).

29. Rives, A., Meier, J., Sercu, T., et al. Biological structure and function emerge from scaling unsupervised learning to 250 million protein sequences. Proceedings of the National Academy of Sciences 118. 10.1073/pnas.2016239118 (2021).

30. Lin, Z., Akin, H., Rao, R., et al. Evolutionary-scale prediction of atomic-level protein structure with a language model. Science 379, 1123–1130. 10.1126/science.ade2574 (2023).

31. Capriotti, E., Fariselli, P. & Casadio, R. I-Mutant2.0: predicting stability changes upon mutation from the protein sequence or structure. Nucleic Acids Research 33, W306–W310. 10.1093/nar/gki375 (2005).

32. Chen, C.-W., Lin, J. & Chu, Y.-W. iStable: off-the-shelf predictor integration for predicting protein stability changes. BMC Bioinformatics 14. 10.1186/1471-2105-14-s2-s5 (2013).

33. Dehouck, Y., Grosfils, A., Folch, B., et al. Fast and accurate predictions of protein stability changes upon mutations using statistical potentials and neural networks: PoPMuSiC-2.0. Bioinformatics 25, 2537–2543. 10.1093/bioinformatics/btp445 (2009).

34. Kellogg, E. H., Leaver-Fay, A. & Baker, D. Role of conformational sampling in computing mutation-induced changes in protein structure and stability. Proteins: Structure, Function, and Bioinformatics 79, 830–838. 10.1002/prot.22921 (2010).

35. Li, G., Panday, S. K. & Alexov, E. SAAFEC-SEQ: A Sequence-Based Method for Predicting the Effect of Single Point Mutations on Protein Thermodynamic Stability. International Journal of Molecular Sciences 22, 606. 10.3390/ijms22020606 (2021).

36. Montanucci, L., Capriotti, E., Frank, Y., et al. DDGun: an untrained method for the prediction of protein stability changes upon single and multiple point variations. BMC Bioinformatics 20. 10.1186/s12859-019-2923-1 (2019).

37. Pandurangan, A. P., Ochoa-Montaño, B., Ascher, D. B., et al. SDM: a server for predicting effects of mutations on protein stability. Nucleic Acids Research 45, W229–W235. 10.1093/nar/gkx439 (2017).

38. Pires, D. E. V., Ascher, D. B. & Blundell, T. L. mCSM: predicting the effects of mutations in proteins using graph-based signatures. Bioinformatics 30, 335–342. 10.1093/bioinformatics/btt691 (2013).

39. Pires, D. E. V., Ascher, D. B. & Blundell, T. L. DUET: a server for predicting effects of mutations on protein stability using an integrated computational approach. Nucleic Acids Research 42, W314–W319. 10.1093/nar/gku411 (2014).

40. Rodrigues, C. H., Pires, D. E. & Ascher, D. B. DynaMut: predicting the impact of mutations on protein conformation, flexibility and stability. Nucleic Acids Research 46, W350–W355. 10.1093/nar/gky300 (2018).

41. Schymkowitz, J., Borg, J., Stricher, F., et al. The FoldX web server: an online force field. Nucleic Acids Research 33, W382–W388. 10.1093/nar/gki387 (2005).

42. Worth, C. L., Preissner, R. & Blundell, T. L. SDM–a server for predicting effects of mutations on protein stability and malfunction. Nucleic Acids Research 39, W215–W222. 10.1093/nar/gkr363 (2011).

43. Benevenuta, S., Pancotti, C., Fariselli, P., et al. An antisymmetric neural network to predict free energy changes in protein variants. Journal of Physics D: Applied Physics 54, 245403. 10.1088/1361-6463/abedfb (2021).

44. Cao, H., Wang, J., He, L., et al. DeepDDG: Predicting the Stability Change of Protein Point Mutations Using Neural Networks. Journal of Chemical Information and Modeling 59, 1508–1514. 10.1021/acs.jcim.8b00697 (2019).

45. Li, B., Yang, Y. T., Capra, J. A., et al. Predicting changes in protein thermodynamic stability upon point mutation with deep 3D convolutional neural networks. PLOS Computational Biology 16 (ed Fariselli, P.) e1008291. 10.1371/journal.pcbi.1008291 (2020).

46. Pancotti, C., Benevenuta, S., Repetto, V., et al. A Deep-Learning Sequence-Based Method to Predict Protein Stability Changes Upon Genetic Variations. Genes 12, 911. 10.3390/genes12060911 (2021).

47. Fariselli, P., Martelli, P. L., Savojardo, C., et al. INPS: predicting the impact of non-synonymous variations on protein stability from sequence. Bioinformatics 31, 2816–2821. 10.1093/bioinformatics/btv291 (2015).

48. Capriotti, E., Fariselli, P., Rossi, I., et al. A three-state prediction of single point mutations on protein stability changes. BMC Bioinformatics 9. 10.1186/1471-2105-9-s2-s6 (2008).

49. Chen, Y., Lu, H., Zhang, N., et al. PremPS: Predicting the impact of missense mutations on protein stability. PLOS Computational Biology 16 (ed Keskin, O.) e1008543. 10.1371/journal.pcbi.1008543 (2020).

50. Zhou, Y., Pan, Q., Pires, D. E. V., et al. DDMut: predicting effects of mutations on protein stability using deep learning. Nucleic Acids Research 51, W122–W128. ISSN: 1362-4962. 10.1093/nar/gkad472 (2023).

51. Iqbal, S., Li, F., Akutsu, T., et al. Assessing the performance of computational predictors for estimating protein stability changes upon missense mutations. Briefings in Bioinformatics 22. 10.1093/bib/bbab184 (2021).

52. Pancotti, C., Benevenuta, S., Birolo, G., et al. Predicting protein stability changes upon single-point mutation: a thorough comparison of the available tools on a new dataset. Briefings in Bioinformatics 23. 10.1093/bib/bbab555 (2022).

53. Pucci, F., Schwersensky, M. & Rooman, M. Artificial intelligence challenges for predicting the impact of mutations on protein stability. Current Opinion in Structural Biology 72, 161–168. 10.1016/j.sbi.2021.11.001 (2022).

54. Jia, L., Yarlagadda, R. & Reed, C. C. Structure Based Thermostability Prediction Models for Protein Single Point Mutations with Machine Learning Tools. PLOS ONE 10 (ed Zhang, Y.) e0138022. 10.1371/journal.pone.0138022 (2015).

55. Masso, M. & Vaisman, I. I. Accurate prediction of stability changes in protein mutants by combining machine learning with structure based computational mutagenesis. Bioinformatics 24, 2002–2009. 10.1093/bioinformatics/btn353 (2008).

56. Pucci, F., Bourgeas, R. & Rooman, M. Predicting protein thermal stability changes upon point mutations using statistical potentials: Introducing HoTMuSiC. Scientific Reports 6. 10.1038/srep23257 (2016).

57. Saraboji, K., Gromiha, M. M. & Ponnuswamy, M. N. Average assignment method for predicting the stability of protein mutants. Biopolymers 82, 80–92. 10.1002/bip.20462 (2006).

58. Topham, C. M., Srinivasan, N. & Blundell, T. L. Prediction of the stability of protein mutants based on structural environment-dependent amino acid substitution and propensity tables. Protein Engineering Design and Selection 10, 7–21. 10.1093/protein/10.1.7 (1997).

59. Masso, M. & Vaisman, I. I. AUTO-MUTE 2.0: A Portable Framework with Enhanced Capabilities for Predicting Protein Functional Consequences upon Mutation. Advances in Bioinformatics 2014, 1–7. 10.1155/2014/278385 (2014).

60. Jumper, J., Evans, R., Pritzel, A., et al. Highly accurate protein structure prediction with AlphaFold. Nature 596, 583–589. 10.1038/s41586-021-03819-2 (2021).

61. Esposito, D., Weile, J., Shendure, J., et al. MaveDB: an open-source platform to distribute and interpret data from multiplexed assays of variant effect. Genome Biology 20. 10.1186/s13059-019-1845-6 (2019).

62. Rubin, A. F., Min, J. K., Rollins, N. J., et al. MaveDB v2: a curated community database with over three million variant effects from multiplexed functional assays. bioRxiv. 10.1101/2021.11.29.470445 (2021).

63. Bava, K. A. ProTherm, version 4.0: thermodynamic database for proteins and mutants. Nucleic Acids Research 32, 120D–121. 10.1093/nar/gkh082 (2004).

64. Gromiha, M. M., An, J., Kono, H., et al. ProTherm: Thermodynamic Database for Proteins and Mutants. Nucleic Acids Research 27, 286–288. 10.1093/nar/27.1.286 (1999).

65. Gromiha, M. M. ProTherm, version 2.0: thermodynamic database for proteins and mutants. Nucleic Acids Research 28, 283–285. 10.1093/nar/28.1.283 (2000).

66. Gromiha, M. M., Thermodynamic Database for Proteins and MProThermutants: developments in version 3.0. Nucleic Acids Research 30, 301–302. 10.1093/nar/30.1.301 (2002).

67. Kumar, M. D. S. ProTherm and ProNIT: thermodynamic databases for proteins and protein-nucleic acid interactions. Nucleic Acids Research 34, D204–D206. 10.1093/nar/gkj103 (2006).

68. Sarai, A., Gromiha, M. M., An, J., et al. Thermodynamic databases for proteins and protein-nucleic acid interactions. Biopolymers 61, 121–126. 10.1002/1097-0282(2002)61:2%3C121::aid-bip10077%3E3.0.co;2-1 (2001).

69. Nikam, R., Kulandaisamy, A., Harini, K., et al. ProThermDB: thermodynamic database for proteins and mutants revisited after 15 years. Nucleic Acids Research 49, D420–D424. 10.1093/nar/gkaa1035 (2020).

70. Xavier, J. S., Nguyen, T.-B., Karmarkar, M., et al. ThermoMutDB: a thermodynamic database for missense mutations. Nucleic Acids Research 49, D475–D479. 10.1093/nar/gkaa925 (2020).

71. Nair, P. S. & Vihinen, M. VariBench: A Benchmark Database for Variations. Human Mutation 34, 42–49. eprint: https://onlinelibrary.wiley.com/doi/pdf/10.1002/humu.22204.https://onlinelibrary.wiley.com/doi/abs/10.1002/humu.22204 (2013).

72. Hernández, I. M., Dehouck, Y., Bastolla, U., et al. Predicting protein stability changes upon mutation using a simple orientational potential. Bioinformatics 39, btad011. ISSN: 1367-4811. eprint: https://academic.oup.com/bioinformatics/article-pdf/39/1/btad011/48769997/btad011.pdf. z10.1093/bioinformatics/btad011 (2023).

73. Ingraham, J., Garg, V., Barzilay, R., et al. Generative Models for Graph-Based Protein Design in Advances in Neural Information Processing Systems (eds Wallach, H., Larochelle, H., Beygelzimer, A., et al.) b32 (Curran Associates, Inc., 2019). https://proceedings.neurips.cc/paper_files/paper/2019/file/f3a4ff4839c56a5f460c88cce3666a2b-Paper.pdf.

74. Akdel, M., Pires, D. E. V., Pardo, E. P., et al. A structural biology community assessment of AlphaFold2 applications. Nature Structural & Molecular Biology 29, 1056–1067. 10.1038/s41594-022-00849-w (2022).

75. Blondel, M., Teboul, O., Berthet, Q., et al. Fast Differentiable Sorting and Ranking in INTERNATIONAL CONFERENCE ON MACHINE LEARNING, VOL 119 (eds Daume, H. & Singh, A.) 119. International Conference on Machine Learning (ICML), ELECTR NETWORK, JUL 13-18, 2020 (2020).

76. Pucci, F., Bernaerts, K. V., Kwasigroch, J. M., et al. Quantification of biases in predictions of protein stability changes upon mutations. Bioinformatics 34 (ed Valencia, A.) 3659–3665. ISSN: 1367-4811. 10.1093/bioinformatics/bty348 (2018).

77. Usmanova, D. R., Bogatyreva, N. S., Ariño Bernad, J., et al. Self-consistency test reveals systematic bias in programs for prediction change of stability upon mutation. Bioinformatics 34 (ed Valencia, A.) 3653–3658. ISSN: 1367-4811. 10.1093/bioinformatics/bty340 (2018).

78. Tsuboyama, K., Dauparas, J., Chen, J., et al. Mega-scale experimental analysis of protein folding stability in biology and design. Nature 620, 434–444. ISSN: 1476-4687. 10.1038/s41586-023-06328-6 (2023).

79. Rodrigues, C. H., Pires, D. E. & Ascher, D. B. DynaMut2: Assessing changes in stability and flexibility upon single and multiple point missense mutations. Protein Science 30, 60–69. ISSN: 1469-896X. 10.1002/pro.3942 (2020).

80. Buel, G. R. & Walters, K. J. Can AlphaFold2 predict the impact of missense mutations on structure? Nature Structural & Molecular Biology 29, 1–2. 10.1038/s41594-021-00714-2 (2022).

81. Hu, M., Yuan, F., Yang, K. K., et al. Exploring evolution-aware & -free protein language models as protein function predictors 2022. arXiv: 2206.06583 [q-bio.QM].

82. McBride, J. M., Polev, K., Reinharz, V., et al. AlphaFold2 can predict single-mutation effects on structure and phenotype. bioRxiv. 10.1101/2022.04.14.488301 (2022).

83. Pak, M. A., Markhieva, K. A., Novikova, M. S., et al. Using AlphaFold to predict the impact of single mutations on protein stability and function. PLOS ONE 18 (ed Budisa, N.) e0282689. 10.1371/journal.pone.0282689 (2023).

84. Zhang, Y., Li, P., Pan, F., et al. Applications of AlphaFold beyond Protein Structure Prediction. bioRxiv. eprint:https://www.biorxiv.org/content/early/2021/11/04/2021.11.03.467194.full.pdf.https://www.biorxiv.org/content/early/2021/11/04/2021.11.03.467194 (2021).

